# Genome assemblies and genetic maps highlight chromosome-scale macrosynteny in Atlantic acroporids

**DOI:** 10.1101/2023.12.22.573044

**Authors:** Nicolas S Locatelli, Sheila A Kitchen, Kathryn H Stankiewicz, C Cornelia Osborne, Zoe Dellaert, Holland Elder, Bishoy Kamel, Hanna R Koch, Nicole D Fogarty, Iliana B Baums

**Affiliations:** Department of Biology, The Pennsylvania State University, University Park, PA, USA; Department of Marine Biology, Texas A&M University at Galveston, Galveston, TX, USA; Institute for System Biology, Seattle, WA, USA; Australian Institute of Marine Science, Townsville, Queensland, Australia; Lawrence Berkeley National Laboratory, Joint Genome Institute, Berkeley, CA, USA; Mote Marine Laboratory, Coral Reef Restoration Program, Summerland Key, FL, USA; Department of Biology and Marine Biology, University of North Carolina Wilmington, Wilmington, NC, USA; Helmholtz Institute for Functional Marine Biodiversity (HIFMB), Ammerländer, Heerstraße 231, 26129 Oldenburg, Germany

**Keywords:** Acropora, coral, genome, chromosome, ancestral linkage group, linkage map, recombination rate, heterochiasmy, hermaphrodite

## Abstract

**Background:** Corals belong to the Cnidaria, an early branching phylum of metazoans. Over the course of their long evolutionary history, they have adapted to changing environments, such as rising sea levels and increasing ocean temperatures. While their history speaks to their evolutionary capacity, it is less clear how quickly they may respond to rapid changes. A critical aspect of adaptive capacity is the structure of their genome and the genetic diversity contained within.

**Findings:** Here, we present chromosome-scale genome assemblies and genetic linkage maps of two critically endangered coral species, *Acropora palmata* and *A. cervicornis,* the two extant Atlantic acroporid corals. Genomes of both species were resolved into 14 chromosomes with comparable assembly sizes (*A. palmata*, 287Mb; *A. cervicornis*, 305Mb). Gene content, repeat content, gene collinearity and macrosynteny were largely preserved between the Atlantic acroporids but a 2.5 Mb inversion and 1.4 Mb translocation were detected between two of the chromosome pairs. Macrosynteny and gene collinearity decreased when comparing Atlantic with Pacific acroporids. Paracentric inversions of whole chromosome arms characterized *A. hyacinthus*, specifically. In the larger context of cnidarian evolution, the four acroporids and another scleractinian coral with chromosome-resolved genome assemblies retained six of 21 cnidarian ancestral linkage groups, while also privately sharing numerous ALG fission and fusion events compared to other distantly related cnidarians. Genetic linkage maps were built using a 30K genotyping array with 105 offspring in one family for *A. palmata* and 154 offspring across 16 families for *A. cervicornis*. The *A. palmata* consensus linkage map spans 1,013.42 cM and includes 2,114 informative markers. The *A. cervicornis* consensus map spans 927.36 cM across 4,859 markers. *A. palmata* and *A. cervicornis* exhibited similarly high sex-averaged genome-wide recombination rates (3.53 cM/Mb and 3.04 cM/Mb, respectively) relative to other animals. In our gamete-specific maps, we found pronounced sex-based differences in recombination, known as heterochiasmy, in this simultaneous hermaphrodite, with both species showing recombination rates 2-2.5X higher in eggs compared to sperm.

**Conclusions:** The genomic resources presented here are the first of their kind available for Atlantic coral species. These data sets revealed that adaptive capacity of endangered Atlantic corals is not limited by their recombination rates, with both species exhibiting high recombination rates and heterochiasmy. Nevertheless, the two sister species maintain high levels of macrosynteny and gene collinearity between them. The few large-scale rearrangements detected deserve further study as a potential cause of fertilization barriers between the species. Together, the assemblies and genetic maps presented here now enable genome-wide association studies and discovery of quantitative trait loci; tools that can aid in the conservation of these endangered corals.

## Introduction

Corals are early branching metazoans with a long evolutionary history, first appearing in the fossil record 240 Mya, though phylogenomic analyses suggest the earliest scleractinians emerged around 425 Mya (Stolarski et al. 2011). Several genome assemblies are now complete and reveal substantial similarities between early and late branching metazoans (Simakov et al. 2022), indicating a slow evolutionary rate in the phylum Cnidaria (corals, hydrozoans and jellyfish). Over evolutionary time scales, corals have adapted to changing environments (Budd and Pandolfi 2010), but it is less clear how fast they may adapt to rapid changes. Aspects of adaptive capacity may include the structure of an organism’s genome, the genetic diversity contained within it, and the rate at which genetic diversity is recombined (Campos et al. 2014).

Corals have complex lifestyles: planktonic larvae settle and form sessile adult colonies via polyp budding and branch fragmentation (Baums et al. 2006, Baird et al. 2009, Harrison 2011). During annual broadcast spawning events, adult Atlantic *Acropora* colonies release egg/sperm bundles into the water column where they dissociate (Szmant 1986). Self-fertilization is genet-specific and self-fertilizing genets occur at low frequency in the populations (Baums et al. 2013, Vasquez Kuntz et al. 2022). Larvae develop for a few days in the water column before swimming towards the benthos where they settle and metamorphose (Richmond and Hunter 1990). Once a primary polyp has formed, symbiotic algae in the order Symbiodiniaceae colonize the coral tissue. Adult colonies of Atlantic acroporids most often harbor the species *Symbiodinium ‘fitti’* (Baums et al. 2014). However, recruitment of sexually produced offspring into adult populations of these acroporids is now rare (Harper et al. 2023). Indeed, populations of Atlantic acroporids have declined more than 80% in recent decades throughout the Atlantic and Caribbean due to anthropogenic impacts, infectious diseases, and temperature induced bleaching events (Bruckner and Hill 2009, Dudgeon et al. 2010) leading to their current status as a federally listed threatened species under the US Endangered Species Act.

Genome assemblies are now available from all classes of cnidarians (Holstein 2022). In Anthozoa, the Hexacorallia are represented by dozens of genomes from genera such as *Acropora* (Shinzato et al. 2020; Fuller et al. 2020, Lopez-Natam et al. 2023), *Astrangia* (Stankiewicz et al. 2023), *Exaiptasia* (Baumgarten et al. 2015)*, Nematostella* (Putnam et al. 2007) and the Octocorallia by at least eight genomes from taxa such as *Renilla* (Jiang et al. 2019), *Dendronephthya* (Jeon et al. 2019)*, Xenia* (Hu et al. 2020), and *Heliopora* (Ip et al. 2023). Seven chromosome-resolved assemblies are published for scleractinian corals (Fuller et al. 2020, McKenna et al. 2021, Yu et al. 2022, Thomas et al. 2022, López-Nandam et al. 2023). While most coral species are diploid, other ploidies exist (e.g. *Pocillopora acuta,* Stephens et al. 2022). The ancestral cnidarian chromosome number is seventeen (Zimmermann et al. 2023), but coral genomes generally have fourteen chromosomes (2n = 28; Kenyon 1997) and genome sizes are between 300Mb – 1Gb (eg., Prada et al. 2016, Fuller et al. 2020, Pootakham et al. 2021, Bongaerts et al. 2021). The number of genes is typically 30,000-40,000 but exceptions exist (e.g. *Montipora capitata* and *Porites compressa* in Stephens et al. 2022).

Genetic diversity fuels adaptation by providing targets for selection (e.g. Torda and Quigley 2022, Mathur et al. 2023). Population genetic data indicate that corals are heterozygous and contain substantial genetic diversity over their large geographic ranges (Baums et al. 2012, Voolstra et al. 2023), including the two Atlantic acroporids (Baums et al. 2005, Drury et al. 2016, 2017, Devlin-Durante and Baums 2017, Kitchen et al. 2019, Canty et al. 2021, García-Urueña et al. 2022). Hybridization and introgression among coral populations and species is facilitated by external fertilization of embryos and synchronized mass spawning events (Vollmer and Palumbi 2002, Budd and Pandolfi 2010, Fogarty et al. 2012). Indeed, the two Atlantic acroporids hybridize to form an F1 hybrid and backcrosses of the F1 hybrid into both parent species are observed at a low frequency (Kitchen et al. 2019). Germline mutations account for around 10% of all mutations found in coral tissue (López and Palumbi 2020, López-Nandam et al. 2023) but no direct estimates of mutation rate or selection coefficients that act on those mutations exist.

Recombination allows for the separation of beneficial and detrimental alleles, such that selection may act upon them independently (Felsenstein 1974). However, the role of recombination in adaptive evolution has been the subject of debate. While recombination has the capacity to create new, advantageous genetic combinations, it can also separate existing ones (Otto and Lenormand 2002). Recombination between adaptive loci may impede range expansions prompted by shifts in environmental conditions (Eriksson and Rafajlović 2021). On the other hand, adaptive substitutions are correlated with higher recombination in several systems (Campos et al. 2014, Castellano et al. 2016, Grandaubert et al. 2019). Further, recombination rate varies across individuals, across the genome, and across sexes (Stapley et al. 2017, Sardell and Kirkpatrick 2020). Global patterns of variation between males and females (heterochiasmy) across taxa suggest these differences may be adaptive (Cooney et al. 2021). Heterochiasmy in simultaneously hermaphroditic animals has been found in the limited number of studies published to date (Wang et al. 2009, Li et al. 2012, Theodosiou et al. 2016), and the recombination landscape of different sexes has only been studied in one other coral, *Acropora millepora* (Wang et al. 2009). Here, we focus on studying the recombination landscape of two, critically endangered sister species, *Acropora palmata* and *A. cervicornis* (**Fig. 1**). Both species are simultaneous hermaphrodites that reproduce sexually and asexually via fragmentation (Szmant 1986). Because these are endangered species, understanding their potential to adapt to changes is a pressing issue.

**Figure 1:**
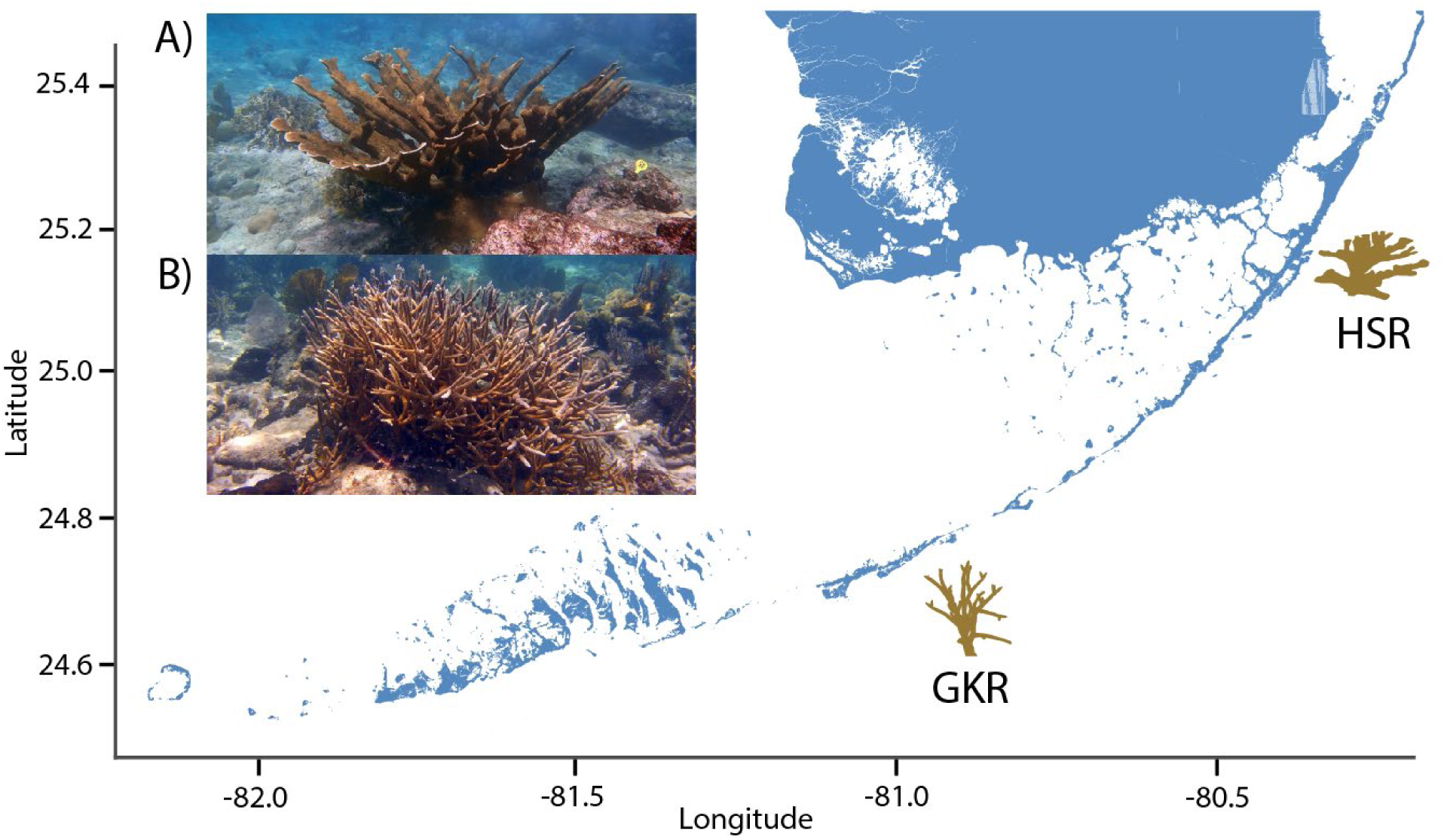
*Acropora palmata* (A) and *A. cervicornis* (B) are dominant reef-building corals of Caribbean and northwestern Atlantic reefs and the only representative species of the genus *Acropora* in the region. Letter notation on the map indicates the geographic origin of *A. palmata* genome genet at Horseshoe Reef (HSR) and *A. cervicornis* genome genet near Grassy Key (GKR). Photos by IBB.

One way to derive recombination rates is by building a genetic linkage map. Linkage maps can be generated from just one cross with many offspring or from few offspring across several families (Rastas 2017). Because one bi-parental coral cross can generate hundreds of offspring, many recombination events can be cataloged among siblings from a few families, or even a single family, and used to order markers along a chromosome. Using a combination of long read, short read, Hi-C chromatin scaffolding, and linkage map anchoring of *de novo* assembled scaffolds, we report chromosome-level genome assemblies of the two Atlantic acroporid species, *Acropora palmata* (Lamarck, 1816) and *A. cervicornis* (Lamarck, 1816). With these assemblies, we compare macrosynteny at the whole genome and gene-level with Pacific acroporids and distant relatives, and characterize the recombination landscapes in these sister species.

## Results and Discussion

### Chromosome-scale genome assemblies of the Atlantic acroporids

To describe the genomic conservation and divergence between the two Atlantic acroporids, we generated chromosome-scale genome assemblies for both species collected from the Florida Keys. For *A. palmata* (genet HS1, STAGdb ID HG0004), we used a hybrid assembly strategy that combined PacBio Sequel II long-reads with Illumina paired-end short reads to obtain an initial assembly with 2,043 scaffolds totaling to 304 Mb and an N50 of 282 kb (N50 is the minimum contig length to cover 50% of the genome). The assembly was further improved with Dovetail Chicago HiRise and Dovetail Hi-C data (**Table S1**). After Hi-C scaffolding, the final 287 Mb haploid assembly was resolved into 14 pseudochromosomes (hereafter referred to as chromosomes, labeled Chr1 - Chr14), a number consistent with the karyotype of *A. palmata* (Devlin-Durante et al. 2016). It is also the most common number of chromosomes shared among acroporids (diploid n=28 in 72% of species surveyed; Kenyon 1997). The *A. palmata* assembly has 406 scaffolds with an N50 of 18.66 Mb (**Fig. 2A** and **Table S2**).

**Figure 2.**
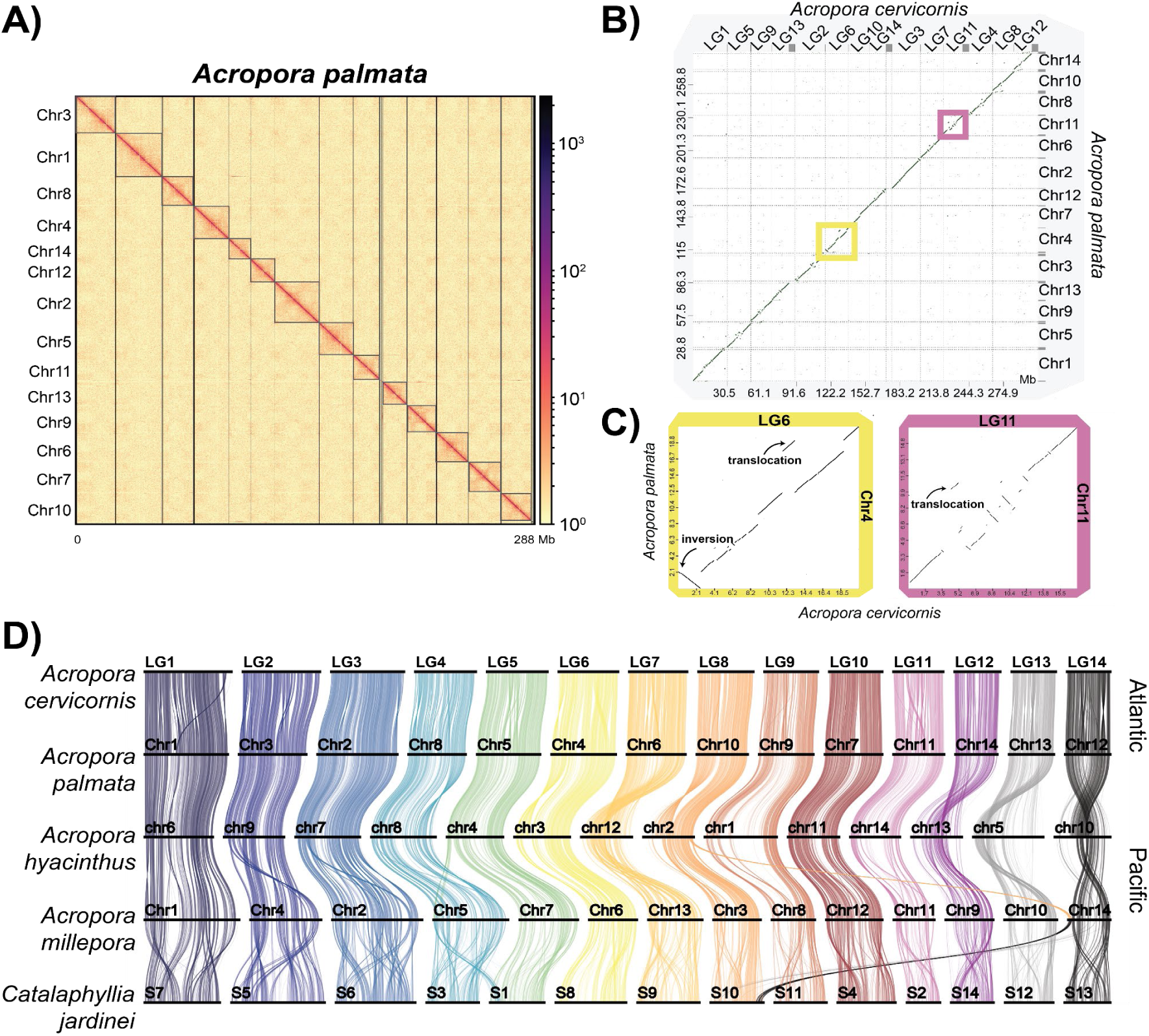
Atlantic acroporid genome assemblies. **(A)** Hi-C contact map of *A. palmata* genome resolved into 14 chromosomes using HiCAssembler (Renschler et al. 2019). **(B)** Dot plot visualization of collinear relationships of the 14 chromosomes/linkage groups between the sister species *A. palmata* (y-axis) and *A. cervicornis* (x-axis) in D-genies web server (Cabanettes and Klopp 2018). The scale on each axis is in megabases (Mb). The points along the diagonal represent collinear genomic regions whereas those dots off the diagonal represent rearrangements (insertions, deletions, inversions and translocations). Yellow and purple boxes highlight two chromosomes, *A. cervicornis* LG6 and LG11, with complex rearrangements. **(C, left)** Comparison of *A. cervicornis* LG6 to *A. palmata* Chr4 reveals an 2.5 Mb inversion and 1.4 Mb translocation. **(C, right)** Complex rearrangements observed between *A. cervicornis* LG11 and *A. palmata* Chr 11, including a 0.765 Mb translocation. **(D)** Ribbon plot of syntenic orthologous genes conserved among scleractinians. The colored vertical links connect orthologous genes to the numbered chromosomes of the five species, represented by horizontal bars. Chromosome fusions or fission are represented by crossing over of the colors that represent each ancestral linkage group. Chromosomal inversions were detected between Atlantic and Pacific acroporids (e.g. *A. cervicorni*s L4, L6, and L12). Chromosomal changes were more numerous between Pacific than Atlantic acroporids. Comparing *A. hyacinthus* Chr 5, 10, 12 and 13 to all other acroporids indicates paracentric inversions of whole chromosome arms in this species.

For *A. cervicornis* (genet GKR, STAGdb ID HG0005), we initially used the same hybrid assembly strategy as for *A. palmata* relying on a combination of PacBio Sequel and Illumina short-read data (**Table S1**). However, due to reduced high molecular weight genomic DNA available at the time, we were unable to size-select our PacBio data as we did for *A. palmata*, yielding shorter read lengths with an average and N_50_ of 3,238 bp and 4,394bp, respectively, compared to 7,126 bp and 10,110 bp in *A. palmata* (**Table S1**). Our first assembly was consequently less contiguous, with 4,382 scaffolds in 318 Mb and an N50 of 162 kb. We next turned to Oxford Nanopore PromethION (ONT) sequencing to generate additional long-read sequences but due to sample quality, the run produced an average read length of 2,366 bp, albeit with much higher overall data yield of 94.4 Gbp. Assembly of the high coverage ONT reads resulted in 6,381 contigs with an N_50_ of 711 Kb. To further resolve the *A. cervicornis* genome, we constructed a linkage map that was used to anchor and orient the ONT scaffolds into 14 linkage groups (LGs). These LGs correspond with high synteny to the Hi-C chromosomes assembled for *A. palmata*. Thus, the *A. cervicornis* LGs can be considered (pseudo)chromosomes. To better distinguish chromosomes for each species, we number the *A. cervicornis* chromosomes here as LG1-LG14. The final 305 Mb assembly was slightly more contiguous than *A. palmata* with an N_50_ of 20.05 Mb.

Recently, a genome assembly of another *A. cervicornis* genotype from the Florida Keys, genet K2 (STAGdb ID HG0582), was published (Selwyn and Vollmer 2023). Using minimap2 (Li 2018) whole genome alignments, we demonstrate that the two assemblies exhibit high sequence homology (**Fig. S1**). Both assemblies are similar in completeness according to BUSCO Metazoa v10 (Manni et al. 2021) assessment with the GKR assembly (this study) showing 93.1% completeness and the K2 assembly showing 92.45% completeness, of which 0.30% and 0.42% are duplicated, respectively (**Table S3**). The assemblies are similar in size, with the GKR assembly being 305 Mb in total length and the K2 assembly 307 Mb. The most notable difference are the gains in contiguity, with a scaffold N50 of 20.051 Mb for the GKR assembly, compared with 2.8 Mb for the K2 assembly. Some K2 contigs are split across multiple linkage groups in the GKR assembly (**Fig. S1**). These regions may reflect novel structural variants between genets within the Florida population of *Acropora cervicornis* or, given the additional linkage scaffolding and misassembly correction used here, may represent a misassembly in the K2 genet.

Our assemblies of the two Atlantic species were on the lower end of the predicted genome sizes from three different k-mer based tools that ranged from 290 to 354 Mb (**Table S4**), and both assemblies are approximately 110 to 180 Mb smaller than genomes of other acroporids species assembled to date (**Table S2**). When comparing estimates of genome completeness using BUSCO Metazoa v10 (Manni et al. 2021), we identified 88% complete genes in *A. palmata*, a reduction that could be due to small local mis-assemblies introduced during the Hi-C scaffolding process or incomplete polishing. Nevertheless, our genome completeness scores are similar to those of other acroporid assemblies (**Table S2**).

### Genomic synteny is largely conserved in the sister species

Whole genome alignments of the two Atlantic acroporid genomes using minimap2 (Li 2018) and nucmer (Marçais et al. 2018) revealed long stretches of collinear regions with interspersed rearrangements across the 14 chromosomes (**Fig. 2B**). As well as similarities, there were differences in physical lengths of chromosomes that resulted in different chromosome number/linkage group assignments for each species (see **Table S5**). For example, the length of the corresponding syntenic chromosome pair of *A. cervicornis* LG2 was 4.87 Mb longer than *A. palmata* Chr3. Overall, we identified 10,532 structural variants totaling 33.02 Mb between the two assemblies using variant calling tools (**Table S6**). An additional 1.4 Mb translocation was detected by whole genome alignment dot plots between *A. cervicornis* LG6 and *A. palmata* chromosome Chr4 (**Fig. 2B and C**). Dot plots also highlighted a large inversion of (2.5 Mb) between the same syntenic chromosome pair (*A. cervicornis* LG6 and *A. palmata* Chr4) and numerous smaller structural variant (SV) types were identified near the middle of *A. cervicornis* LG11 and *A. palmata* Chr11 (**Fig. 2C**), a region that may correspond with the centromere.

Small inversions and translocations should be independently confirmed because marker density of the *A. cervicornis* linkage map was only 16 markers per Mb and contigs containing a single marker cannot be oriented correctly. Lep-Anchor (Rastas 2020) additionally utilizes long-read data to assist in contig orientation where linkage markers are sparse or absent, but in cases where long reads are too short to span repetitive regions, the correct orientation may still not be resolved. Long distance translocations and large-scale inversions may be more immune to these issues. Additionally, because of the presence of unbridged gaps from Hi-C and linkage scaffolding, breakends may not be detected or supported by SV callers despite being detected by alignment dot plots.

The two species discussed here are able to hybridize in nature to form an F1 hybrid, previously referred to as *A. prolifera*, and rare backcrosses of the F1 with both parent species have been documented. However, F2 generations have not been observed in genetic data from wild colonies (Vollmer and Palumbi 2002, Kitchen et al. 2019). Given the paucity of later generation hybrids (backcrosses and F2s), the hybrids may undergo hybrid breakdown resulting in non-viable or less fit offspring. It is therefore assumed that some genetic mechanism, like differing genomic architectures, exists that represses reproduction between the parental species (Vollmer and Palumbi 2002). For example, large structural variants can cause misalignment during F1 meiosis or death in F2 offspring due to the loss of gene copies required for survival (Zhang et al. 2021). Such structural variants cause F2 sterility in interspecies hybrids of *Drosophila* (Masly et al. 2006), as well as F2 lethality in wild strains of *Arabidopsis* (Bikard et al. 2009). Although whole genome alignments between *A. palmata* and *A. cervicornis* demonstrate high levels of macrosynteny and conserved gene collinearity, some regions do exhibit large scale rearrangements (e.g., 2.5Mb inversion on LG6/Chr4, **Fig. 2B** and **C, Table S6**). Structural variants may be acting as a barrier to backcross and F2 offspring formation in the F1 hybrid adults, and represent candidates for future studies of hybrid breakdown in this system.

### Genome architecture and gene content across cnidarians

To predict gene models for each assembly, we used a combination of transcriptomic data and *ab inito* tools resulting in 31,827 and 34,013 genes in *A. palmata* and *A. cervicornis*, respectively (**Table S2**). Combining our gene models with those of other acroporids with chromosome-resolved assemblies, we identified collinear (shared loci with the same arrangement on a given chromosome) and macrosyntenic (shared loci not necessarily in the same arrangement on a given chromosome) gene arrangements (**Fig. 2D and Fig. S2A**). In accordance with the high degree of synteny at the whole genome level, 15,873 out of 17,243 one-to-one orthologs between *A. palmata* and *A. cervicornis* retained their collinearity (**Fig. S2**). The number of orthologs that shared ordinal positions between *A. cervicornis* chromosomes and *A. hyacinthus* or *A. millepora* was 12,603 out of 13,000 and 12,075 out of 14,738, respectively. We found that the architecture of some chromosomes was largely unchanged at this scale of observation (e.g. *A. cervicornis* LG1 across acroporids, **Fig. 2D**). Thus, over their 52 - 119 million years (Mya) of history (Shinzato et al. 2021), acroporids have retained conserved syntenic gene order to a high degree.

Nevertheless, several translocations and inversions were evident. Within the acroporids, interchromosomal translocations were observed in *A. millepora* with 85 genes of *A. cervicornis* LG8 located on Chr 14 of *A. millepora* and 132 genes of *A. cervicornis* LG5 located on *A. millepora* Chr 5 (**Fig. 2D**). Paracentric inversions of whole chromosome arms likely led to the *A. hyacinthus* Chrs 5, 10, 12 and 13 arrangements (**Fig. 2D and Fig. S2A**). In agreement with Ying et al. 2018 and Shinzato et al. 2021, collinear relationships declined with phylogenetic distance from the acroporid lineage (**Fig. 2D, Fig. S2 and Fig. S3**). For example, comparison of the acroporids, members of the complex clade of corals, with the coral *Cataphyllia jardinei*, which belongs to the robust coral clade, show macrosyntenic continuity within the 14 chromosomes (**Fig S2A**) but gene collinearity was mostly lost (**Fig 2D**). While only a small sample size is available for comparison, the maintenance of chromosomal arrangements across deeply diverged coral lineages that split in the Devonian–Carboniferous, approximately 332–357 Mya (Quattrini et al. 2020), is surprising. Macrosyntenic patterns gradually degraded and chromosome numbers varied as we compared acroporids to more divergent species from Actiniaria, Octocorallia and Medusozoa (**Fig. S3**).

Ancestral chromosomal fusions and rearrangements within the coral lineage were detected by mapping previously inferred ancestral linkage groups (ALGs) shared among sponges, cnidarians and bilaterians against our genomes (Simakov et al. 2022). We note changes in ancestral ALGs in the discussion below with fusions represented by the letter “x” (**Table S7**). Of the 21 cnidarian specific ALG arrangements, six (*A1a, Ea, J1xQa, A1bxB3, NxA2,* and *B1xB2*) were largely intact within the scleractinians (acroporids and *Catalaphyllia*), represented by LG7, LG13, LG12, LG11, LG14 and LG5 in *A. cervicornis* (**Fig. S2B** and **Table S7**). Interestingly, ALG *Qb* was lost from all cnidarian species surveyed here, with the exception of the jellyfish *Cassiopea xamachana* that largely retains the ancestral cnidarian ALG structure (**Table S7** and **Fig. S3**). We identified seven cases of ALG fusions and one example of centric insertion within one of the acroporid chromosomes, represented by *A. cervicornis* LG10 (**Fig. S2B** and **Table S7**). *A. millepora* is the only acroporid species where a portion of ALG *G* fused with *L*. This fusion event in *A. millepora* presents an interesting target for further studies in light of the variable hybridization potential among species within the genus.

Expanding beyond the species with chromosome-resolved assemblies, we compared orthologous gene families, also known as orthogroups, shared among diverse cnidarian taxa, including representatives of the Hexacorallia and Octocorallia within Anthozoa and Hydrozoa and Scyphozoa within Medusozoa (**Table S8**). We identified 2,601 conserved orthogroups among all cnidarians (**Fig. 3**). There are 159 unique orthogroups in Anthozoa enriched in the process angiogenesis (GO:0001525, *p.adjust*=0.049) and 44 unique Scleractinia orthogroups enriched in growth factor binding (GO:0019838, *p.adjus*t= 0.009), cell adhesion molecule binding (GO:0050839, *p.adjust* = 0.035) and D-inositol-3-phosphate glycosyltransferase activity (GO:0102710, *p.adjust*= 0.008). We further found 42 and 142 unique orthogroups in acroporids and Atlantic acroporids, respectively (**Fig. 3**). Similar to a prior study (Shinzato et al. 2021), the acroporid-specific groups included gene families involved in coral calcification (galaxin, matrix shell protein and skeletal organic matrix protein) and host-microbe interactions (prosaposin and toll-like receptor). Only 39 of the 142 orthogroups shared between the Atlantic species were annotated, 12 of which were predicted as transposable elements, suggesting numerous coding genes and/or repetitive element copies arose after gene flow stopped between the Atlantic and Pacific acroporids, approximately 2.8 Mya (van Oppen et al. 2001, O’Dea et al. 2016). Notable genes with lineage-specific duplications include a gene involved with sperm function (OG0022455: cation channel sperm-associated protein 3), two involved in DNA replication (OG0022558: Serine/threonine-protein kinase Nek2 and OG0022391: replication protein A 70 kDa DNA-binding subunit C) and one in development (OG0022485: paired box protein).

**Figure 3.**
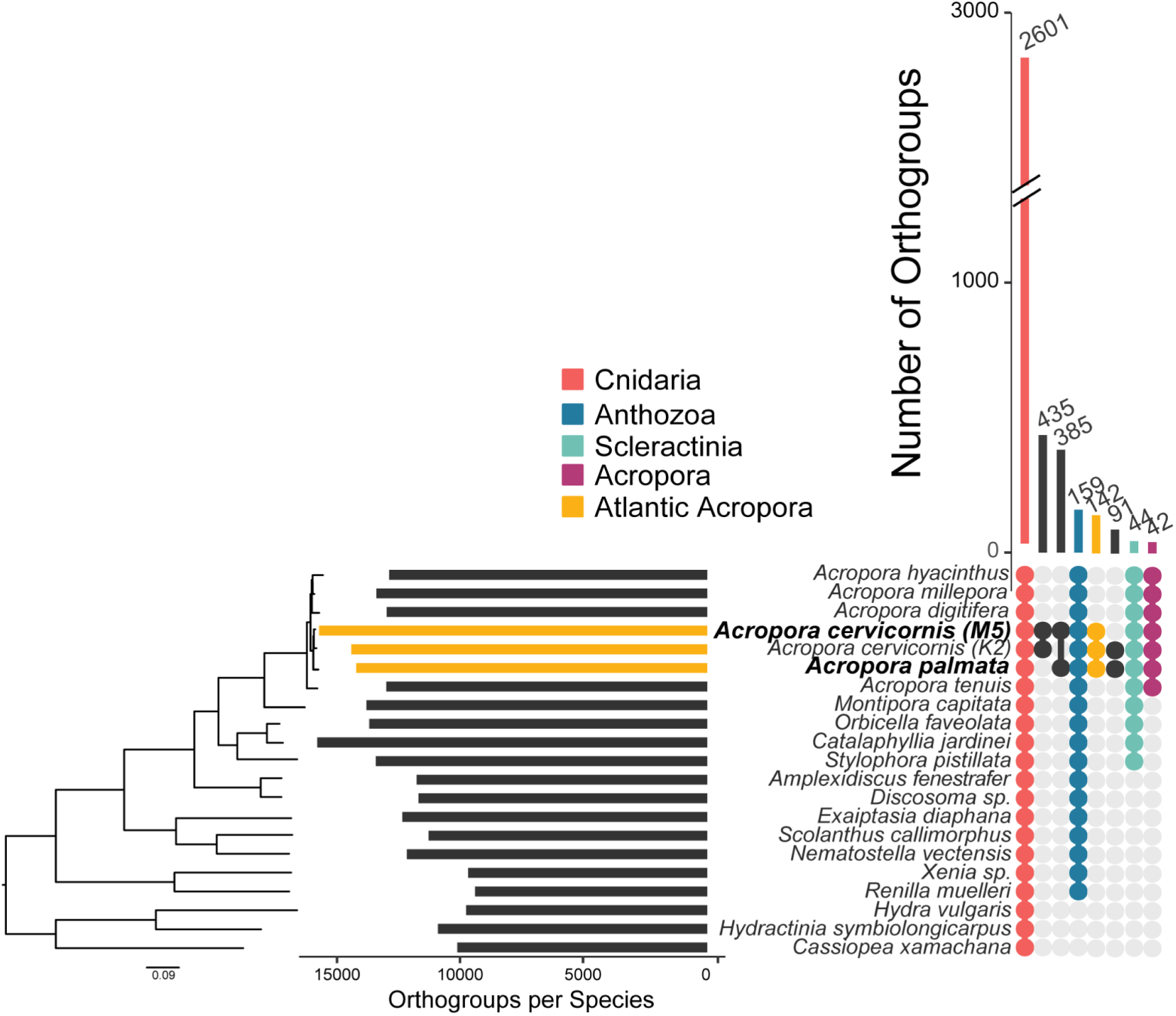
Conservation of gene content among cnidarians. UpSet plot displaying the number of shared orthologous groups among taxa. The colored or black circles below the vertical bar chart indicate those species that belong to each group. Groups highlighted include Cnidaria (red), Anthozoa (blue), Scleractinia (green), *Acropora* (purple) and Atlantic (Caribbean) *Acropora* (yellow). On the left, the bar chart represents the total number of orthologous groups identified in each taxon. Taxon labels in bold were assembled in this study. The species tree constructed from 1,011 orthogroups was inferred by STAG and rooted by STRIDE in OrthoFinder v2.5.2 (Emms and Kelly 2019).

### Repetitive content is comparable among acroporids

Repetitive DNA plays a significant role in the size, organization and architecture of eukaryotic genomes (Feschotte and Pritham 2007). To analyze transposable element (TE) content among the acroporid genome assemblies, we constructed species-specific repeat libraries for each assembly using a genome-guided approach with RepeatModeler (Flynn et al. 2020). To ensure that only bona fide repeats were included in our comparisons, we filtered out putative genes using a sequence similarity approach against the NCBI protein database or *A. digitifera* gene models. Despite their smaller genome sizes, we found the TE content of the Atlantic acroporids, 16.69% in *A. palmata* and 18.91% in *A. cervicornis* (**Table S9**), was similar to other acroporid species whose TE content ranged from 13.57% in *A. digitifera* to 19.62% in *A. tenuis* (**Fig. 4A**). It should be noted that our predicted TE content for all species is lower than previous estimates of 40% to 45% for acroporids using a different TE identification method (Shinzato et al. 2021), suggesting that we may have underestimated total repeat content. Using dnaPipeTE, an assembly-free method based on the Illumina short-reads, total TE content was estimated to be 37.11% for *A. palmata* and 35.54% for *A. cervicornis* (**Fig. S4**), supporting our estimates with RepeatMasker were low. The distribution of repeats assigned to each class differed slightly between methods and studies, reflecting the limitations of using a single tool for TE identification and annotation (Rodriguez and Arkhipova 2023). Regardless, the genome size differences between the Atlantic and Pacific species cannot be attributed to a reduction or expansion in genomic TE content in the respective lineages.

**Figure 4.**
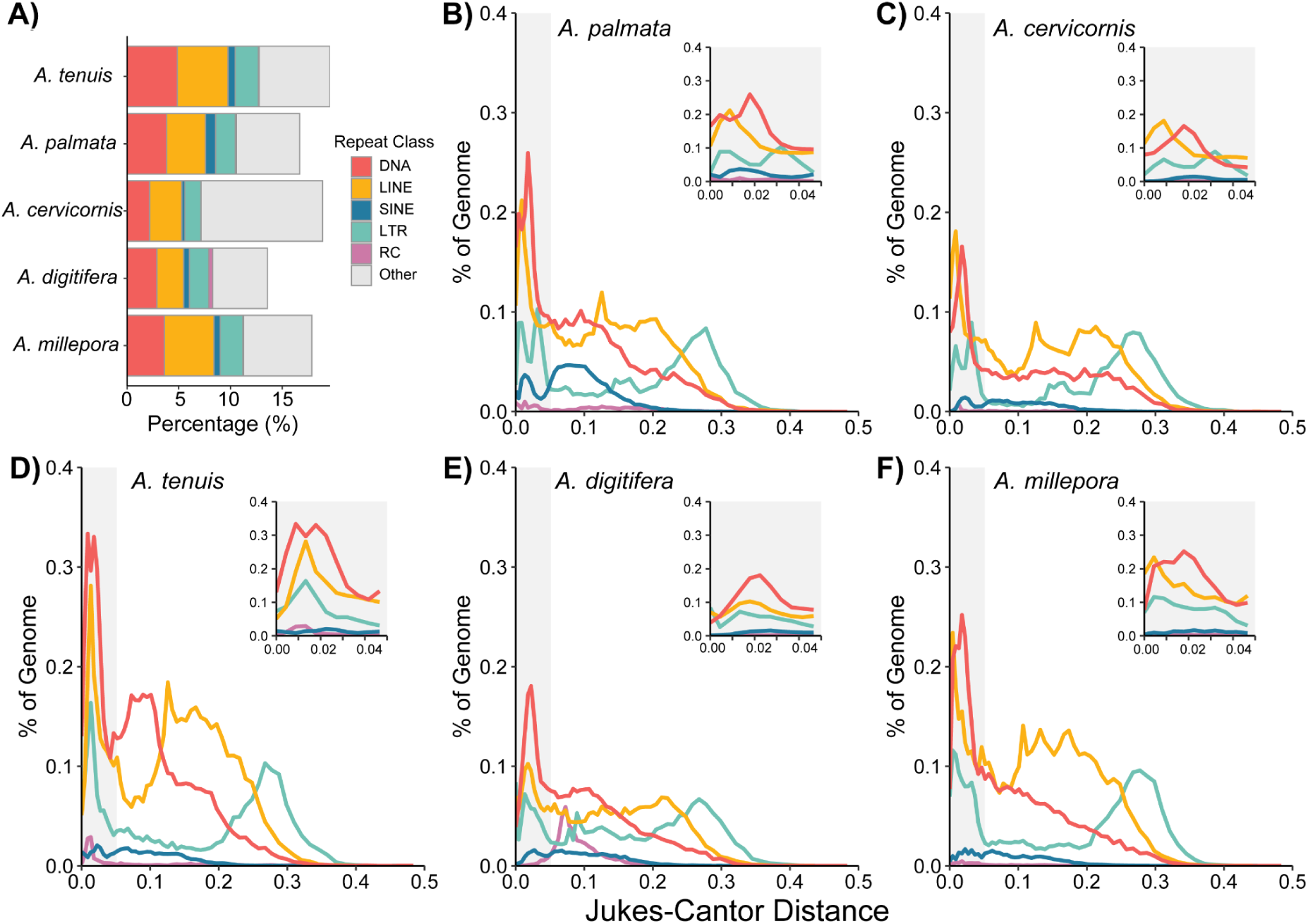
Comparison of repetitive DNA among acroporid taxa. **(A)** Percentage of the genome attributed to the main transposable element classes [DNA transposons, long interspersed nuclear element (LINE), short interspersed nuclear element (SINE), long terminal repeat (LTR), rolling circle (RC) and other (satellites, simple repeats, and unclassified)] for each acroporid taxon. **(B-F)** Repeat landscapes of all transposable element classes except “other” for *A. palmata* **(B)**, *A. cervicornis* **(C)**, *A. tenuis* **(D)**, *A. digitifera* **(E)** and *A. millepora* **(F)**. The percentage of genome coverage (y-axis) of each repeat is shown relative to the Jukes-Cantor genetic distance observed between a given repetitive element and its respective consensus sequence. Individual repetitive elements were then summarized by their repeat class. The more recent repetitive element copies have lower Jukes-Cantor distance on the left side of the x-axis. The inset plot in each panel focuses on recent repeat insertions at a Jukes-Cantor distance below 0.05 (gray shaded region in full plot).

The dominant TEs were shared among the species we surveyed across methods. These TEs belong to DNA transposons superfamilies Tc/Mariner and hAT, long interspersed nuclear element (LINE) retrotransposon family Penelope and long terminal repeat (LTR) family Gypsy (**Table S9**). The transposable activity of each repeat class was compared across species to determine if TE accumulation differed over their evolutionary past (**Fig. 4B-F**). Each species experienced a recent burst of DNA, LINE and LTR copies in their genomes, as evidenced by the increased genomic coverage of those classes with zero to very small genetic distances (**Fig. 4B-F inset plots**). Within the recent TE expansion, the Atlantic acroporids and *A. millepora* have a bimodal distribution of LTR transpositions, specifically those within the retrotransposon family Gypsy. Overall, however, few species-specific patterns emerged in the repeat landscapes of acroporids.

### Genetic Maps

#### Acropora palmata genetic linkage map

In total, we assigned 2,114 informative markers to 14 linkage groups (LGs), representing the 14 chromosomes of the *A. palmata* genome, with an average marker distance of 0.48 cM and a consensus map length of 1,013.42 cM (**Table 1**). The gamete-specific maps varied in length, with a higher female map length (1,460.68 cM) than the male map length (583.19 cM). Marker number and density varied across chromosomes with the highest number of markers associated with Chr1 (318) and the lowest in Chr14 (82). Examination of the genetic position (cM) against the physical position (Mb) of each marker in the genome showed high agreement between the linkage map and the genome assembly. In the female map, the LG length ranged from 79.95 cM to 148.29 cM. In contrast, in the male map, the LG length ranged from 27.67 cM to 59.69 cM. The consensus map LGs ranged from 53.63 cM to 100.30 cM. In all 14 LGs, the female length was longer than the male length (**Table 1**). Analysis of gamete-specific linkage maps in *A. palmata* revealed sexual dimorphism with respect to genome-wide and chromosome-level recombination rate (heterochiasmy). The genome-wide average recombination rate was higher in the female (5.49 cM/Mb) than in the male (2.19 cM/Mb) (**Table 1**). The highest average recombination rate (7.00 cM/Mb) was in the female map associated with Chr11. The lowest average recombination rate (1.55 cM/Mb) was in the male map associated with Chr2. In all 14 chromosomes, the female recombination rate was higher than the male rate.

**Table 1:**
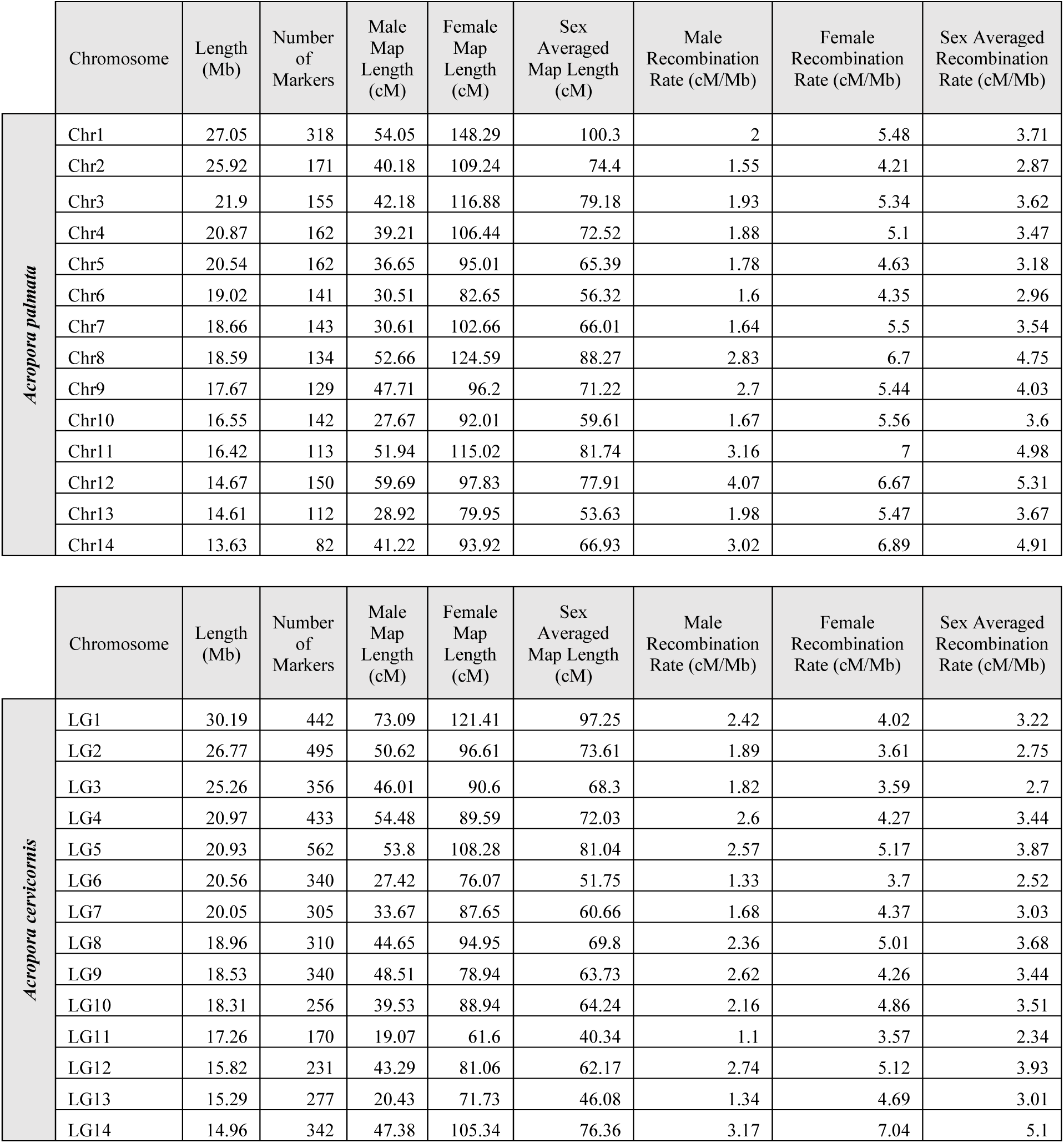
Physical lengths, map length, and average recombination rates per chromosome for male, female, and sex-averaged maps of *Acropora palmata* and *A. cervicornis*. Mb = megabases, cM = centimorgan.

#### Acropora cervicornis genetic linkage map

The *A. cervicornis* linkage map was constructed with more offspring (154) from 16 families, and thus a greater number of informative markers were utilized in generating a consensus linkage map. In total, 4,859 markers were assigned to 14 linkage groups (LGs), with an average marker distance of 0.19 cM and a consensus map length of 927.36 cM (**Table 1**). The gamete-specific maps varied in length, with a higher female map length (1,252.78 cM) than the male map length (601.93 cM). Marker number and density varied across LGs with the highest number of markers in the linkage group associated with LG5 (562) and the lowest in LG11 (170). In the female map, the LG length ranged from 61.60 cM to 121.41 cM. In contrast, in the male map, the LG length ranged from 19.07 cM to 54.48 cM. The consensus map LGs ranged from 40.34 cM to 97.25 cM. As in *A. palmata,* for all 14 *A. cervicornis* LGs, the female length was longer than the male length (**Table 1**) and pronounced heterochiasmy was detected. The genome-wide average recombination rate was higher in the female (4.41 cM/Mb) than in the male (2.12 cM/Mb) (**Table 1**). The highest average recombination rate (7.04 cM/Mb) was in the female map associated with LG14. The lowest average recombination rate (1.10 cM/Mb) was in the male map associated with LG11. In all 14 linkage groups, the female recombination rate was higher than the male rate (**Table 1**).

### Interspecies comparisons between Atlantic acroporids

Linkage maps were largely concordant between species, with recombination rates and centromere positions similar across taxa, as highlighted in **Fig. 5** and **Fig. S5**. However, some notable differences in map length and recombination rates were present. One homologous chromosome pair (LG11/Chr11) exhibited large differences in map length in which the linkage map for *A. palmata* was almost twice as long as the map for *A. cervicornis*, despite similar physical size (115cM vs. 61.6cM in the female map). In both species, heterochiasmy was pronounced, with female recombination rates roughly two times as high as male rates in *A. cervicornis* and roughly 2.5 times as high in *A. palmata*. Heterochiasmy in *A. palmata* and *A. cervicornis* was among the most pronounced estimates observed in plants or animals (Brandvain and Coop 2012). Generally, recombination rates were higher in *A. palmata*, potentially due to differences in overall assembly length. The k-mer estimated genome sizes were similar (333 Mb in *A. palmata* and 331 Mb in *A. cervicornis*, **Table S3**) but assembly sizes were more variable, with *A. palmata* being 287 Mb and *A. cervicornis* being 305 Mb. This would result in increased genome-wide *A. palmata* recombination rates simply due to assembly size. However, regardless of assembly sizes, genetic map lengths are greater in *A. palmata* (consensus map length 1013 cM) than in *A. cervicornis* (927 cM). Based on repeat density and local recombination rates (i.e. regions with elevated repeat content and suppressed recombination, as described in Hartley and O’Neill 2019 and Schreiber et al. 2022), all chromosomes in both species appear to be metacentric or submetacentric (**Fig. 5** and **Fig. S5**), like in the Pacific acroporid, *Acropora pruinosa* (Taguchi et al. 2020). Centromeric regions appear to be associated with long interspersed nuclear element (LINE) repeats, as shown by the prominent peaks in LINE content.

**Figure 5:**
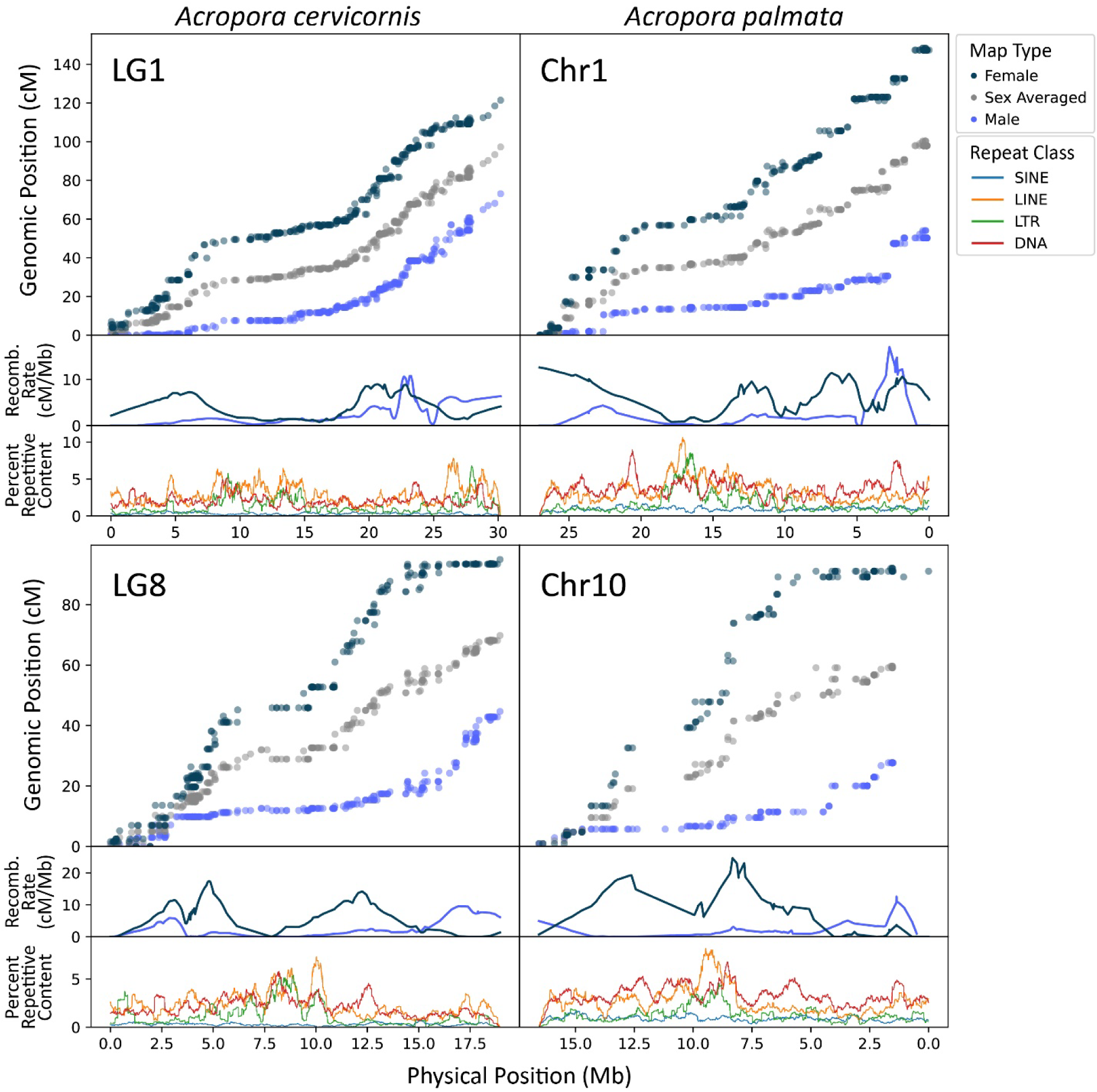
Genetic and recombination maps for two homologous pairs of chromosomes of *Acropora cervicornis* and *A. palmata*. LG1 and Chr1 are homologous, as well as LG8 and Chr10. Percent SINE, LINE, LTR, and DNA repeats show putative centromere positions. Repeat content was calculated in 500Kb sliding windows with 5Kb steps. Note: *A. palmata* Chr1 and Chr10 x-axes indicating physical position are inverted due to the assembled sequence being reverse of the homologous chromosome in *A. cervicornis*.

Within chromosomes, both species exhibit commonly observed local recombination landscapes (e.g., higher local recombination rates in females across whole chromosomes or higher recombination in males near telomeres; Sardell and Kirkpatrick 2020). Twelve out of fourteen chromosomes exhibit recombination landscapes where local female rates are generally higher than male rates throughout the chromosome. Female maps exhibit marked declines in recombination around the presumed centromere while males show low, chromosome-wide recombination. However, in two cases, male local recombination rates are higher than female rates at one end of the chromosome, in telomeric regions (LG9/Chr9, LG8/Chr10, **Fig. 5** and **Fig. S5**). Higher telomeric recombination rates in males have been documented in other animal systems (e.g., humans; Bhérer et al. 2017, frogs; Brelsford et al. 2016, geese; Torgasheva and Borodin 2017). The asymmetry of centromere position is associated with telomeric recombination in male stickleback (Sardell et al. 2018), with elevated telomeric recombination only occurring in long arms in acrocentric chromosomes while short arms exhibit near complete suppression of recombination. In both cases of elevated telomeric recombination (LG8/Chr10 and LG9/Chr9), we observed elevated male telomeric recombination only on the long arm (centromere position inferred by elevated LINE repeat content and locally reduced recombination).

The local and genome-wide recombination rates calculated from the genetic linkage map for *A. palmata* and *A. cervicornis* provide novel insights into the recombination landscape of corals. The density of markers in this resource now opens the possibility for quantitative trait locus (QTL) analyses as well as more precise haplotype imputation and genetic association studies in these species (**Fig. 5**). QTL mapping allows for the identification of loci that have consistent, predictable effects on phenotype across individuals. In plants, this is frequently used to assist with breeding programs (Kulwal, 2018). As populations of many corals have rapidly declined (Hughes et al., 2017), such a tool could assist in the design of restoration approaches. Additionally, phasing and imputation softwares commonly used in genome-wide association studies (GWAS) such as BEAGLE (Browning et al. 2018), GLIMPSE2 (Rubinacci et al. 2023), and SHAPEIT (Delaneau et al. 2019) take into account recombination rates across chromosomes to more accurately statistically phase and impute data. The generation of these assemblies and genetic maps now enables complex genetic association studies not previously possible in these threatened non-model organisms. With these data, we have also demonstrated the application of the Acropora SNP array (Kitchen et al., 2020) as a successful genotyping method for the generation of a genetic linkage map, which provides a cost-effective means for creating additional maps for the F1 hybrid of *A. palmata* and *A. cervicornis*.

### Interspecies comparisons among acroporids and other organisms

We present the second and third published coral genetic maps to date, and so interspecific comparisons were finally possible among coral species, *A. palmata*, *A. cervicornis*, and *Acropora millepora* (Wang et al. 2009), as well as between other non-coral taxa.

Comparing recombination among *Acropora palmata*, *A. cervicornis* and *A. millepora* revealed many similarities among the Atlantic (*A. palmata, A. cervicornis*) and Pacific (*A. millepora*) corals. Though characterization of local recombination was not possible in *A. millepora* due to the lack of an assembled genome at the time, Wang et al. (2009) provided important insights into coral recombination by using map lengths to discover heterochiasmy in this species. Both *A. palmata* and *A. cervicornis* (this study) and *A. millepora* linkage maps indicated the presence of 14 chromosomes, consistent with the *A. palmata* karyotype and karyotypes present in many other scleractinian corals (Kenyon 1997, Devlin-Durante et al. 2016, Kawakami et al. 2022). In *A. palmata, A. cervicornis,* and *A. millepora*, the female map length was longer than the male length. While in *Acropora millepora*, higher recombination in the female map was driven by only a subset of linkage groups (Wang et al. 2009), we find that in *A. palmata* the pattern is consistent across all chromosomes (**Table 1**, **Fig. 5**, and **Fig. S5**).

The average genome-wide recombination rate for *A. palmata* and *A. cervicornis* in the female (5.089 and 4.107 cM/Mb, respectively), male (2.03 and 1.97 cM/Mb), and consensus maps (3.53 and 3.03 cM/Mb) is relatively high for an animal and echoes the findings in *A. millepora* (Wang et al. 2009). Across currently available taxa, the recombination rates in *Acropora* were similar to insects, crustaceans, and fish, but were higher than the averages for groups such as birds, amphibians, reptiles, and mammals (Stapley et al. 2017). This could indicate a rapid response to selection in acroporids because the proportion of substitutions fixed by adaptive evolution is positively correlated with recombination rate (Cavassim et al. 2021). Future work comparing recombination rates across coral populations and taxa would be valuable in clarifying the evolutionary consequences of these patterns.

## Conclusions

The genomic resources presented here revealed that the adaptive capacity of endangered Atlantic corals is not hindered by their recombination rates, as both species exhibit high, genome-wide recombination rates with prominent heterochiasmy between sexes in these simultaneous hermaphrodites. The two Atlantic species exhibit remarkable levels of macrosynteny and gene collinearity with one another, and with Pacific species, especially considering the >50Mya history of the genus. Our assemblies suggest that, like many scleractinians, the Atlantic acroporid genome consists of 14 chromosomes; a derived state compared to the last common ancestor of the Cnidaria which is proposed to have had 17 chromosomes (Zimmermann et al. 2023). Over evolutionary time, coral species merge and separate across the tropical oceans with sea-level changes (Veron 1995) and mutation-drift equilibrium is thus seldom, if ever, achieved (Benzie 1999). The conserved number of haploid chromosomes among many, but not all, of the acroporids is 14 (2n=28, Kenyon 1997) and the high level of macrosynteny across the *Acropora* genus may enable these syngameons described above. In the Pacific, it has been suggested that hybridization acts as an evolutionary force driving speciation (Willis et al. 2006, Richards et al. 2008). However, in the Atlantic *Acropora*, hybridization between the two sister species yields a functionally sterile F1 hybrid (Vollmer and Palumbi 2002) despite the high levels of macrosynteny and gene collinearity of their genomes. Together, the chromosome-scale assemblies and genetic maps we present here are the first detailed look at the genomic landscapes of these critically endangered species. The availability of these genomic resources helps facilitate genome-wide association studies and discovery of quantitative trait loci which can aid in the conservation of endangered corals.

## Material and Methods

### Sample collection and sequencing

Adult coral tissue collected from the *Acropora cervicornis* genet GKR collected near Grassy Key (24.711783° N, 80.945966° W) and reared at the Coral Restoration Foundation Tavernier Nursery (CRF, 24.9822° N, 80.4363° W) and the *A. palmata* genet HS1 from Horseshoe Reef (25.1399° N, 80.2946° W) were described previously (Table S1; Kitchen et al. 2019). High molecular weight genomic DNA (gDNA) was isolated from each coral tissue sample using the Qiagen DNeasy kit (Qiagen, Valencia, CA) with slight modifications described previously (dx.doi.org/10.17504/protocols.io.bgjqjumw). Paired-end 250 bp sequencing libraries (avg. insert size 550 nt) were constructed from 1.8-2 µg gDNA with the TruSeq DNA PCR-Free kit (Illumina, San Diego, CA) and sequenced on the Illumina HiSeq 2500 by the Genomics Core Facility at Pennsylvania State University. Additionally, coral tissue from *A. palmata* HS1 was collected by CRF in January of 2018, snap-frozen in liquid nitrogen and sent directly to Dovetail Genomics for DNA extraction followed by Chicago and Hi-C library preparation.

For the PacBio libraries, gamete bundles of *A. cervicornis* GKR (spawned 2015 and August 22, 2016 at the CRF nursery) and *A. palmata* HS1 (spawned August 20, 2016 at Horseshoe Reef) were collected during the annual coral spawn. Once the gamete bundles broke apart, sperm was separated from the eggs using a 100 μm filter, and concentrated and washed with 0.2 μm filtered seawater through three rounds of centrifugation at 2,000 x g for 5 min at room temperature. The *A. cervicornis* sperm samples from 2015 were brought to a final concentration of 3 x 10^7^ cells ml^-1^ after the addition of Cell Suspension Buffer and 2% agarose using the Bio-Rad CHEF Genomic DNA Plug Kits (Bio-Rad, Hercules, CA). Genomic DNA plugs were processed according to the manufacturer’s protocol and stored at 4 °C. The genomic DNA was extracted from the plugs in two ways, either using the QIAquick Gel Extraction kit (Qiagen) or by soaking the plugs overnight in 100 ul nuclease-free water at 4 °C followed by 1 h at −80 °C and recovered at 23,000 x g. Sperm samples of both species from 2016 were stored as 1 ml aliquots of concentrated sperm in 100% non-denatured ethanol at −20 °C until extraction. Genomic DNA was extracted using Nucleon Phytopure DNA extraction kit (Cytiva, Marlborough, MA) with the addition of RNase treatment and increased incubation time of 3 to 4 h at 65 °C during the cell lysis step. Genomic DNA elutions were combined and concentrated using the AMPure bead clean-up (final gDNA= 2 μg for *A. cervicornis* and 10 μg for *A. palmata*). Given the different final gDNA concentrations, PacBio libraries were prepared using a 20kb size-selection protocol for *A. palmata* and a low input, no size selection protocol for *A. cervicornis*. Both libraries were sequenced on Sequel II by the Genomics Core Facility at Pennsylvania State University.

Because the initial *A. cervicornis* assembly exhibited low contiguity, an additional assembly was generated using Oxford Nanopore (ONT) long-read sequencing data. For the *A. cervicornis* ONT DNA library, coral tissue from the GKR genotype preserved in ethanol was provided by the Coral Restoration Foundation in 2021 and stored at −20°C until extraction. Genomic DNA was extracted using the Qiagen MagAttract HMW DNA kit (MD, USA) following the manufacturer’s protocol. To further purify the gDNA, a salt-ethanol precipitation was performed. Briefly, 0.1 volumes of 3M NaAc (pH 5.2) were added to the DNA elution, followed by 3 volumes of 100% ethanol. The sample was centrifuged at approximately 20,000 x g for 1 h at 4 °C. The supernatant was then removed and the pellet was washed twice with cold 75% EtOH. The dried pellet was resuspended in Buffer AE (Qiagen, MD, USA) and sequenced on a Oxford Nanopore PromethION by the University of Wisconsin Biotechnology Center.

### K-mer genome size estimation

We removed low-quality bases (Phred score below 25) and adaptors from Illumina reads, discarding reads shorter than 50 bp, with cutadapt v1.6 (Martin 2011). Prior to genome assembly, 119-mer counting was performed on trimmed reads from each sample using jellyfish v2.2.10 (Marcais and Kingsford 2012) for the purpose of haploid genome size estimation. We tested k-mer 119 because it was identified as the best k-mer for *de novo* genome assembly from contamination filtered reads by KmerGenie v1.7048 (Chikhi and Medvedev 2014) after testing a range of k-mers from 21 to 121. K-mer frequency histograms were analyzed using the *GenomeScope2* web portal (Ranallo-Benavidez et al. 2020) and findGSE (Sun et al. 2018), which use a negative binomial and skew distribution model, respectively.

### Contamination filtering of Illumina short read data

DNA extractions on the adult tissue used for Illumina sequencing were composed of the coral host and its associated microbial partners (algal symbionts and other microbes). To remove non-coral reads, we applied a modified series of filtering steps that compares sequence homology and GC content similar to process in BlobToolKit (Kumar et al. 2013, Challis et al. 2020) and described previously for *A. cervicornis* by (Reich et al. 2021). Adaptor trimmed reads were initially assembled into contigs with SOAPdenovo2 v0.4 (parameters -K 95 –R) (Luo et al. 2012). The contigs were compared to the genomes of the coral *Acropora digitifera* (NCBI: GCF_000222465.1; Shinzato et al. 2011), the symbiont *Breviolum minutum* (OIST: symbB.v1.0.genome.fa; Shoguchi et al. 2013), and the NCBI nucleotide database (nt) using megablast (evalue 1e^-5^ threshold) (Altschul et al. 1997). Contigs with higher sequence similarity to non-cnidarians in the nt database were combined to make a local contamination database. Adaptor trimmed reads were then aligned with Bowtie2 v. 2.2.9 (parameters –q –fast; (Langmead and Salzberg 2012) sequentially against the *A. digitifera* mitochondria (NBCI: KF448535.1), three concatenated Symbiodiniaceae genomes (*Symbiodinium microadriaticum, B. minutum, Fugacium kawagutii*; Shoguchi et al. 2013, Lin et al. 2015, Aranda et al. 2016, respectively) and the contamination database. Unaligned reads were extracted and used for short-read genome assembly described below.

### Hybrid genome assembly of A. cervicornis and A. palmata

The trimmed and filtered short reads were assembled with SoapDeNovo-127mer v2.04 (Luo et al. 2012) using different k-mers for each species, *A. palmata* K= 99 and *A. cervicornis* K= 95. Contigs were filtered for additional symbiont contamination using megablast against the three Symbiodiniaceae genome assemblies described above. A surprising number of symbiont contigs, roughly 500,000 in each species assembly, were present despite our read contamination filtering (Reich et al. 2021). The non-symbiont contigs were then assembled with PacBio long reads using the hybrid method DBG2OLC (Ye et al. 2016, k=17 MinLen=500 AdaptiveTh=0.001 KmerCovTh=2 MinOverlap=20). PacBio reads were also assembled separately with Canu v1.5 (Koren et al. 2017, genomeSize=400m correctedErrorRate=0.075 minReadLength=500). The two assemblies (hybrid and PacBio only) were then combined using QuickMerge v0.2 (Chakraborty et al. 2016, *A. palmata* = -hco 5.0 -c 1.5 -l 55000 -ml 1000; *A. cervicornis* = -hco 5.0 -c 1.5 -l 99500 -ml 1000) with the hybrid assembly as the reference and PacBio assembly as the query. Additional contig extension was performed with FinisherSC v2.1 (Lam et al. 2015). Lastly, the assemblies were polished using Pilon v1.22 (Walker et al. 2014).

### Hi-C scaffolding of hybrid Acropora palmata assembly

Our hybrid assembly of *A. palmata* was submitted to Dovetail Genomics for Hi-C analysis. They combined their proprietary HiRise scaffolding and Hi-C analysis (**Table S1**), but the assembly was still far from chromosome resolved (441 scaffolds, N_50_ = 6.8 Mb, and L_50_ = 16 scaffolds). In an effort to further improve the *A. palmata* genome assembly, we mapped the Hi-C paired-end reads separately back onto the Dovetail Genomics assembly with bwa-mem v 0.7.17 (Li 2013) with the mapping parameters -A1 -B4 -E50 -L0. We then followed the steps outlined by HiCExplorer v2.1.1 to create and correct a Hi-C contact matrix using default settings with a lower bin correction threshold of −1.5 (Ramírez et al. 2018). This indicated there were more short range (< 20kb) than long range (> 20kb) contacts in the matrix. The corrected matrix was then used by HiCAssembler v1.1.1 (Renschler et al. 2019) to further orient the scaffolds into pseudochromosomes with a minimum scaffold length set to 300,000 bp, a bin size of 15,000 and two iterations.

### Nanopore assembly Acropora cervicornis

PromethION data was trimmed and filtered with *Porechop* (Wick et al. 2017), resulting in a total of 94 Gb across 39.91 M reads of usable ONT data. With trimmed ONT data, *metaFlye* (Kolmogorov et al. 2020) was used to perform a long-read only metagenome assembly. Following the initial *metaFlye* assembly, which includes a long-read polishing step, the assembly was further polished in one round using *hypo* (Kundu et al. 2019). Illumina short read data from the GKR genet described above was trimmed using *TrimGalore* (Krueger et al. 2021), and mapped to the preliminary assembly with *bwa-mem* (Li 2013) prior to use with *hypo*. ONT reads were then mapped to the assembly using *minimap2* (Li 2018) and BAM files were sorted using *samtools* (Danecek et al. 2021). Using *blastn* (Camacho et al. 2009), assemblies were searched against a custom database comprised of NCBI’s *ref_euk_rep_genomes, ref_prok_rep_genomes, ref_viroids_rep_genomes*, and *ref_viruses_rep_genomes* databases combined with dinoflagellate and *Chlorella* genomes (Shoguchi et al. 2013, 2018, 2021, Hamada et al. 2018, Beedessee et al. 2020). Using the mapping and *blastn* hits files, *blobtools* (Laetsch and Blaxter 2017) was used to identify and isolate cnidarian contigs. *Purge_dups* (Guan et al. 2020) was utilized to identify and remove any remaining putative haplotigs in the respective assembly.

### Linkage map construction

A full-sibling family was generated through a controlled cross between two *Acropora palmata* genets. Spawn was collected from two genets during the August 2018 spawning season in Curacao. Once egg-sperm bundles had broken apart, gametes were separated and eggs were washed to remove any remaining self-sperm. The sperm from the genet designated as the sire was used to fertilize washed eggs from the genet designated as the dam. The resulting larvae were reared to 96 hours post-fertilization in filtered seawater before preservation in individual 1.5ml PCR tubes with 96% ethanol. A total of 105 full-sibling offspring were used in the construction of the genetic linkage map. Three to four polyps of each spawning parent were collected using coral cutters and preserved in 96% ethanol. For *Acropora cervicornis*, coral recruits from 16 families reared in a previous study until they first branched (Koch et al. 2022). Samples of these recruits were preserved in 95% ethanol in 1.5 mL Eppendorf tubes and immediately placed into a - 80° freezer until extraction.

For *Acropora palmata* larval offspring, high molecular weight DNA extractions followed the methods in Kitchen et al. (2020). Each larva was incubated in 12 μl of lysis solution (10.8 μl Buffer TL, 1 μl of Proteinase K, and 0.2 μl of 100 mg/ml RNAse A, all reagents from Omega BioTek) for 20 min at 55 °C. Next, 38 μl of Buffer TL and 50 μl of phenol/chloroform/isoamyl alcohol solution (25:24:1) was added to each sample and gently rocked for approximately 2 mins. After centrifuging each sample for 10 mins at 20,000g, the top aqueous phase was removed and placed in a new tube. 50 μl of chloroform:isoamyl alcohol (24:1) was added to each sample and gently rocked for 2 minutes. Samples were centrifuged again at 10,000 rpm for 5 mins and the top aqueous phase was again removed and placed into a new tube. The DNA was precipitated with 1.5x volume of room-temperature isopropanol, 1/10 volume of 3M sodium acetate (pH=5.2) and 1 μl of glycogen (5 mg/ml) for 10 mins at room temperature. Samples were then centrifuged at 20,000g for 20 mins and washed with 70% ice-cold ethanol. All supernatant was removed and pellets were dried under a hood for approximately 30 mins. Pellets were re-suspended in 30 μl of low TE buffer (10 mM Tris-HCl and 0.1 mM EDTA). Parental tissue was extracted using Qiagen DNeasy kit (Qiagen, Valencia, CA) following the modified protocol described in Kitchen et al. (2020) and eluted in 100 μl of nuclease-free water.

Extracted samples were genotyped using the Applied Biosystems Axiom Coral Genotyping Array—550962 (Thermo Fisher, Santa Clarita, CA, USA). The raw data were analyzed using the Axiom ‘Best Practices Workflow’ (BPW) with default settings (sample Dish QC ≥ 0.82, plate QC call rate ≥97; SNP call-rate cutoff ≥97; percentage of passing samples ≥ 95). The resulting genotyping files were converted to variant caller format (VCF) using the bcftools plugin affy2vcf (https://github.com/freeseek/gtc2vcf) and filtered to represent only the recommended probeset identified by the Axiom BPW.

*A. cervicornis* recruits were sampled from the base and DNA was extracted by Eurofins BioDiagnostics (WI, U.S.A) using LGC (Hoddesdon, UK) Sbeadex Animal DNA Purification Kits. Samples were run on two plates of the Applied Biosystems Axiom Coral Genotyping Array. *A. cervicornis* cross data was processed in the same manner as *A. palmata*, using the Axiom workflow and subsetting single nucleotide variants to only include recommended probes.

*Acropora palmata* and *A. cervicornis* linkage analysis was carried out using *Lep-MAP3* (Rastas 2017) using the wrapper pipeline *LepWrap* (available at https://github.com/pdimens/LepWrap, Dimens 2022). Markers were first filtered for deviation from Mendelian inheritance and missing data via the *Lep-MAP3* module ParentCall2. For *A. cervicornis*, the flag halfSibs=1 was added to ParentCall2 to account for shared parentage among crosses. Recombination informative markers (here defined as those that were heterozygous in at least one parent) were next filtered using the Filtering2 module with a data tolerance of 0.0001. The remaining markers were assigned to 14 linkage groups (LGs) using an LG minimal size limit set to 5 markers using the module SeperateChromosomes2 and a logarithm of odds (LOD) score of 11 in *A. palmata* and 5 in *A. cervicornis*. For *A. palmata*, an informativeMask value of “123” was used and for *A. cervicornis* multi-family data, an informativeMask of “12” was used. Unassigned markers were iteratively added to existing LGs using a LOD limit of 2 and a LOD difference of 2. Markers were next ordered using the Kosambi mapping function as implemented in the module OrderMarkers2 with the identical limit set to 0.005, usePhysical=1 0.1, 100 merge iterations, 3 phasing iterations, and the hyperPhaser parameter used to improve marker phasing. To remove markers at map edges that may erroneously inflate the map length, the last 10% of markers were trimmed if they fell more than 5% of the total centimorgan (cM) span away from the next nearest marker. After trimming, marker order was evaluated with a second round of OrderMarkers2 using the same parameters as previously described. Both paternal and maternal maps were generated and the option sexAverage = 1 was applied to include a sex-averaged consensus map. Average marker distance was calculated as the size of the linkage map in cM divided by the number of markers. As global orientation of a linkage group is arbitrary in *Lep-MAP3*, marker order was flipped for LGs in which the start of the genetic map (0 cM) corresponded to the end, rather than to the start of the physical map (the position 0 bp) of a given scaffold. To generate cleaned Marey maps, MareyMap Online (Siberchicot et al. 2017) was used to remove aberrant markers and generate smoothed recombination maps using 2-degree polynomial LOESS estimation with a span of 0.25.

### Linkage scaffolding of A. cervicornis Nanopore assembly

For *A. cervicornis*, no Hi-C data was available. As such, the *A. cervicornis* assembly was scaffolded using *Lep-Anchor* (Rastas 2020) with the linkage map generated by *Lep-MAP3* (Rastas 2017). To assist in orientation of contigs with markers, as well as placements of contigs without markers, *minimap2* v2.24 (Li 2018) was used to generate a PAF file using the ONT data. *Lep-Anchor* was run via *LepWrap* and utilized default *Lep-Anchor* arguments, with the exception of setting the expected number of linkage groups to 14. Additionally, *LepWrap* implements the edge-trimming scripts for *Lep-Anchor* as was described above for *Lep-MAP3*.

### Repeat identification, masking and divergence analysis

For both assemblies, repetitive sequences were predicted with RepeatModeler v 1.0.11 (Flynn et al. 2020), filtered for genuine genes based on blast similarity to the NCBI nr database or *Acropora digitifera* protein sequences (e-value ≤ 1e^-5^), combined with the *Acropora* TE consensus sequences in Repbase (n=149), annotated separately against the invertebrate repeat database in CENSOR v4.2.29 (Jurka et al. 1996) for “unknown” TEs, and soft masked using RepeatMasker v 4.0.7 (Smit et al. n.d.). We also ran the above series of steps on the genome assemblies of *A. digitifera*, *A. tenuis* and *A. millepora* to ensure comparable repeat estimates. The summary table for each species was generated using the *buildSummary.pl* utility script, and TE accumulation was calculated as the Kimura substitution level corrected for CpG content from the respective consensus sequence produced using the *calcDivergence. pl* and *createRepeatLandscape.pl* utility scripts in RepeatMasker. Kimura distance was converted to Jukes-Cantor distance using the formula JC = −3/4*log(1 − 4\**d*/3), where *d* is the distance estimated by RepeatMasker. Assembly-free repeat identification, annotation and quantification was performed on 25% of the adapter-trimmed Illumina short-read data of each Atlantic species using dnaPipeTE v1.3.1 (Goubert et al. 2015).

### Gene prediction and annotation

For the *A. palmata* assembly, we used a combination of *ab inito* (GeneMark-ES v4.32; Brůna et al. 2020) and reference-based tools (BRAKER v2.0; Brůna et al. 2021, PASA v2.1.0; Haas et al. 2008), and exonerate v2.2.0; Slater and Birney 2005) for gene prediction as previously described (Brückner et al. 2022). For BRAKER, RNAseq data produced on the Roche 454 GS FLX Titanium system was obtained from NCBI Bioproject PRJNA67695 (Polato et al. 2011) and mapped to the assembly using STARlong v2.5.3a (Dobin et al. 2013) due to the average read lengths being greater than 300 bp. Gene models with read coverage greater than or equal to 90% were assigned as “BRAKER_HiQ’’ predictions. The assembled *A. palmata* transcriptome from Polato et al. 2011 was used as the input for PASA. Homology-based gene predictions were made with exonerate against all eukaryotic sequences in the UniProt database (n=186,759), keeping predictions with at least 80% percent coverage. Gene predictions were combined with EVidenceModeler (Haas et al. 2008). We also predicted tRNA sequences using tRNAscan_SE v1.3.1 (Lowe and Eddy 1997). The predicted genes were searched against the NCBI nr, UniProt Swiss-Prot and Treml databases, and KEGG Automated Annotation Server. Blast-based searches were filtered by the top hit (e-value 1e-5 threshold). GO annotations were extracted from UniProt of NCBI databases. Genes were also compared to OrthoDB v10.1 (Kriventseva et al. 2019). Gene annotation was assigned based on the e-value score < 1e-10 first to Swiss-Prot followed by Trembl and then NCBI. If no sequence homology was recovered, then the gene was annotated as a “hypothetical protein”. Gene predictions from the hybrid assembly were lifted over to the final Hi-C assembly using the UCSC liftOver process (Kuhn et al. 2013). We also used homology-based prediction tool GeMoMA v1.6.1 (Keilwagen et al. 2019) to map the *A. palmata* gene models to the Hi-C assembly. Liftover and GeMoMa predictions were combined with EVidenceModeler for the final gene set.

The original PacBio *A. cervicornis* assembly was annotated in a similar manner to *A. palmata*. However, the original assembly is superseded here by the ONT-based assembly. The ONT *A. cervicornis LepWrap-* scaffolded assembly was annotated using *funannotate v1.8.13* (Palmer and Stajich 2020) with RNAseq data obtained from four BioProjects available on NCBI SRA at the time of assembly (PRJNA222758, PRJNA423227, PRJNA529713, and PRJNA911752). All RNAseq data was adapter- and quality-trimmed using *TrimGalore* (Krueger et al. 2021). Briefly, *funannotate train* was run with a *–max_intronlen* of 100000. *Funannotate train* is a wrapper that utilizes *Trinity* (*68*) and *PASA* (*69*) for transcript assembly. Upon completion of training, *funannotate predict* was run to generate initial gene predictions using the arguments *–repeats2evm*, *--organism other*, *–max_intronlen 100000,* and *--repeat_filter none*. Additional transcript evidence from three sources (initial *A. cervicornis* annotation described above, transcripts from the Selwyn and Vollmer 2023 assembly, and the Osborne 2023 transcriptome) was provided to *funannotate predict* using the *–transcript_evidence* argument. *Funannotate predict* is a wrapper intended to separately run *AUGUSTUS* (Stanke et al. 2006) and *GeneMark* (Brůna et al. 2020) for gene prediction and *EvidenceModeler* (Haas et al. 2008) to combine gene models. *Funannotate update* was run to update annotations to be in compliance with NCBI formatting. For problematic gene models, *funannotate fix* was run to drop problematic IDs from the annotations. Finally, functional annotation was performed using *funannotate annotate* which annotates proteins using *PFAM* (Bateman et al. 2004), *InterPro* (Hunter et al. 2009), *EggNog* (Huerta-Cepas et al. 2019), *UniProtKB* (Boutet et al. 2016), *MEROPS* (Rawlings et al. 2009), *CAZyme* (Huang et al. 2018), and *GO* (Harris et al. 2004).

### Whole genome alignments

Genome assemblies of *A. palmata*, *A. cervicornis* GKR genet and *A. cervicornis* K2 genet were aligned using minimap2 (Li 2018) with “asm5” setting for whole genome alignments, and the *nucmer* command within the mummer v4.0 package (Marçais et al. 2018) with a minimum exact match length of 100 bp (-l 100), minimum cluster length of 500 (-c 500) and using all anchor positions (--maxmatch). The PAF alignments from minimap2 were plotted using both R package pafr v0.0.2 (David Winter 2020) and dotplotly (https://github.com/tpoorten/dotPlotly). The delta alignments from *nucmer* were visualized using the D-Genies web server (Cabanettes and Klopp 2018). Structural variants (insertions, deletions, tandem duplications and contractions, inversions and translocations) were identified from the whole genome alignments of *A. palmata* and *A. cervicornis* GKR genet using three tools: assemblytics (Nattestad and Schatz 2016), MUM&Co (O’Donnell and Fischer 2020), and SVIM-asm (Heller and Vingron 2021). Only MUM&Co and SVIM-asm were able to detect inversions and translocations (**Table S6**).

### Orthologous gene identification and macrosynteny analysis

Genome completeness of each acroporid assembly was assessed using BUSCO v4.1.1 with the Metazoa odb10 orthologous gene set (n=954 orthologues, Manni et al. 2021). To discover shared and unique gene families in *A. cervicornis* and *A. palmata* in relation to other species, *OrthoFinder* v2.5.2 (Emms and Kelly 2019) was run on the predicted proteins of each species listed in **Table S7.** The species tree was constructed with STAG and rooted by STRIDE in OrthoFinder v2.5.2 (Emms and Kelly 2019). A presence/absence table of orthogroups, or sets of genes descended from a single gene in the last common ancestor of all the species being considered, was used to generate an UpSet venn diagram made with UpSetR v1.4.0 (Conway et al. 2017). We extracted shared orthogroups from select taxonomic groupings highlighted in **Fig. 4** and performed GO enrichment tests with clusterProfiler v4.4.4 (Wu et al. 2021) using a custom database for *A. palmata* created with AnnotationForge v1.38.0 (Marc Carlson 2017).

Macrosyntenic patterns across the species with chromosome-resolved genome assemblies was assessed with Oxford Dot Plots (ODP, Schultz et al. 2023), specifically mapping on the inferred ancestral linkage groups (ALGs) of sponge, cnidarian and bilaterians recently identified (Simakov et al. 2022). ODP runs an all-vs-all blast akin to OrthoFinder with diamond v2.0.15 (Buchfink et al. 2015) and identifies conserved syntenic gene arrangements between two genomes. Dot plots and ribbon diagrams were generated by ODP with default settings and restricting plotted scaffold length of 2 Mb to visualize conserved syntenic blocks across closely related or more distant taxa.

## Author contributions

Locatelli NS performed research, analyzed data, assembled A. cervicornis genome, constructed A. cervicornis linkage map, wrote the manuscript

Kitchen SA performed research, analyzed data, assembled A. palmata genome, assembled first draft A. cervicornis genome, wrote the manuscript

Stankiewicz KH performed research, analyzed data, generated A. palmata linkage map, wrote the manuscript

Dellaert Z performed research, edited manuscript

Elder H, analyzed data, edited manuscript

Kamel BH, analyzed data, edited manuscript

Koch HR, provided data, edited manuscript

Osborne CC, performed research, edited manuscript

Fogarty N, provided funding, edited manuscript

Baums IB, performed research, provided funding, wrote the paper, designed study

## Acknowledgements

The authors wish to thank the Caribbean Research and Management of Biodiversity (CARMABI) research station, the Coral Restoration Foundation (CRF), the Pennsylvania State University Huck Institute of the Life Sciences Genomics Core Facility, University of Wisconsin Biotechnology Center, Valerie Chamberland, Greg von Kustner, Meghann Devlin-Durante, Hannah G Reich, Kate L Vasquez-Kuntz, Cody Engelsma, Marina Villoch, Erich Bartels, Trinity L Conn, Erinn M Muller, Samuel Vohsen, and Abigail Clark for their invaluable contributions to the project.

## Permits

Research was conducted under the following permits:

CRF-2016-020-2016 spawning trip, GKR sperm collection only

CRF-2017-009 and NOAA #FKNMS-2011-159-A4-2017 spawning collection for the hybrid crosses

CRF-2017-012 and NOAA #FKNMS-2011-159-A4 – Horseshoe *A. palmata* fragment for the chromosome-level genome assembly at Dovetail

NOAA #FKNMS-2019-012 A1, A2, A3, and A4, SAJ-2019-04431-(SP-GGM), PER00414 – GKR ethanol-preserved tissue for Nanopore sequencing

NOAA #FKNMS-2015-163 A2, A3 - 2020 spawning of *A. cervicornis* and production of linkage map offspring

## Data availability

Genome assemblies and annotations available upon request. NCBI accessions for assemblies and raw sequencing data will be provided upon final publication. Annotations, auxiliary files, and code associated with each genome assembly will be available on GitHub for final publication.

