## Supplemental Figures and Tables for "Genome assemblies and genetic maps highlight chromosome-scale macrosynteny in Atlantic acroporids"

### Supplementary Data

#### Figures

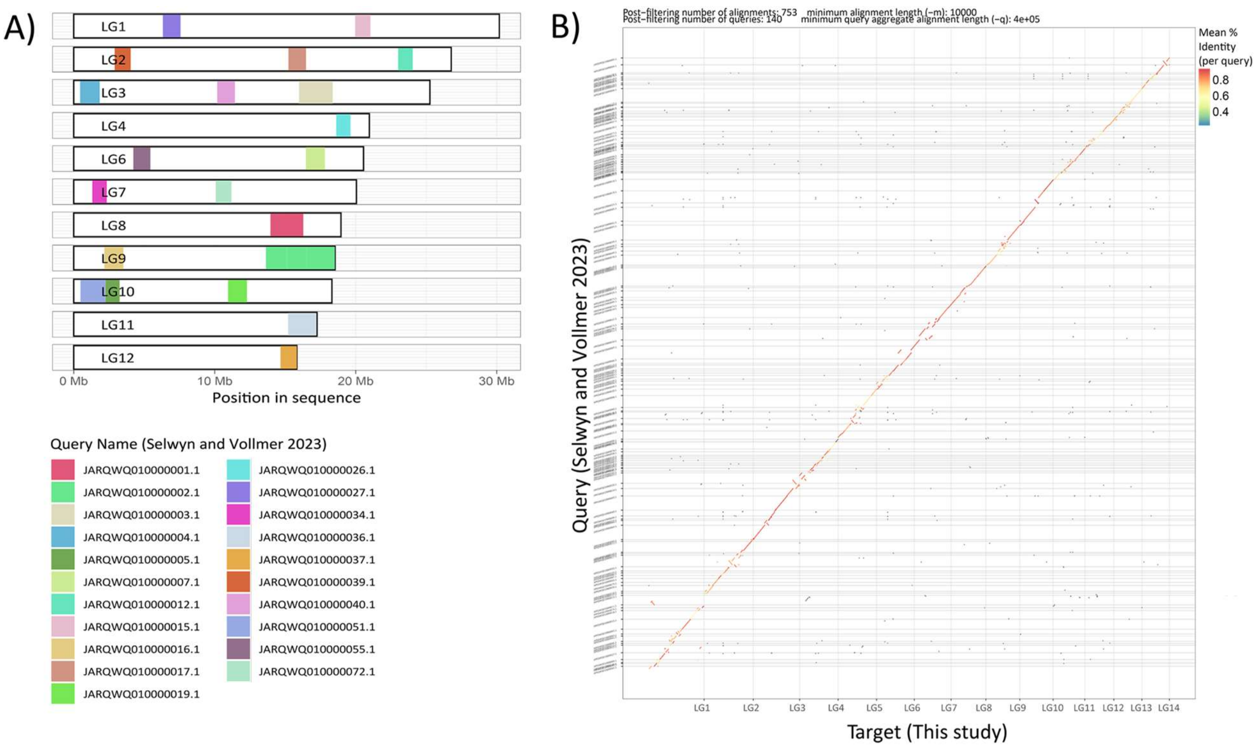


**Figure S1. Alignments of the *Acropora cervicornis* genet K2 assembly (STAGdb ID HG0582, Selwyn and Vollmer 2023) with the genet GKR assembly (STAGdb ID HG0005, this study).** A) Assembly-assembly minimap2 alignments >1Mbp and their chromosome locations, plotted with pafr. B) Dot plots comparing the HG0582 assembly with the 14 HG0005 chromosomes (plotted with dotploty, https://github.com/tpoorten/dotPlotly). Contigs are largely concordant with the exception of one large HG0582 contig that was split across HG0005 chromosomes.

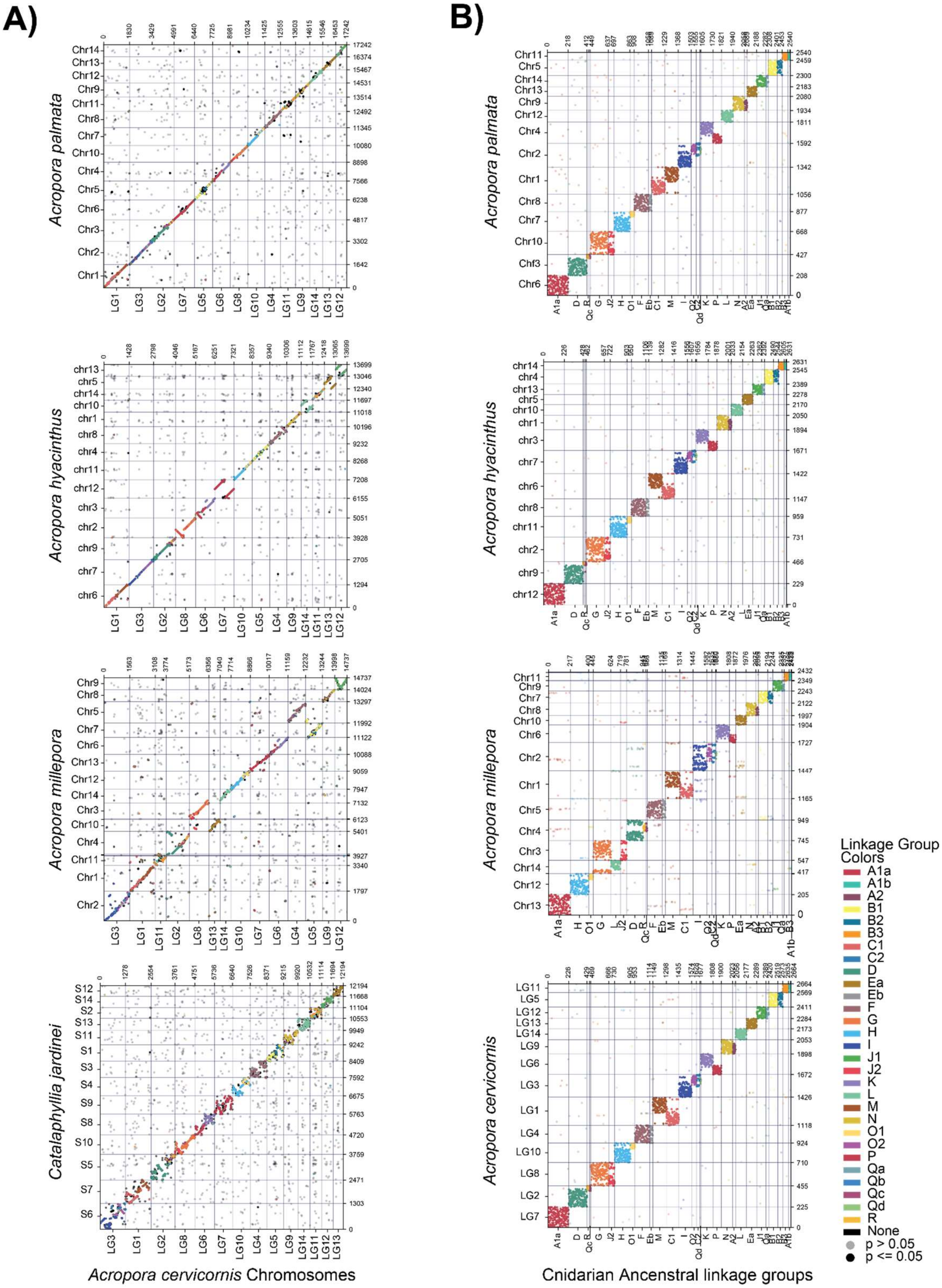


**Figure S2. Oxford dot plots of scleractinian chromosomes against *A. cervicornis* chromosomes (A) and of acroporids chromosomes against 29 ancestral linkage groups (ALGs) of last common ancestor of sponges, cnidarians and bilaterians (Simakov et al. 2022) (B).**

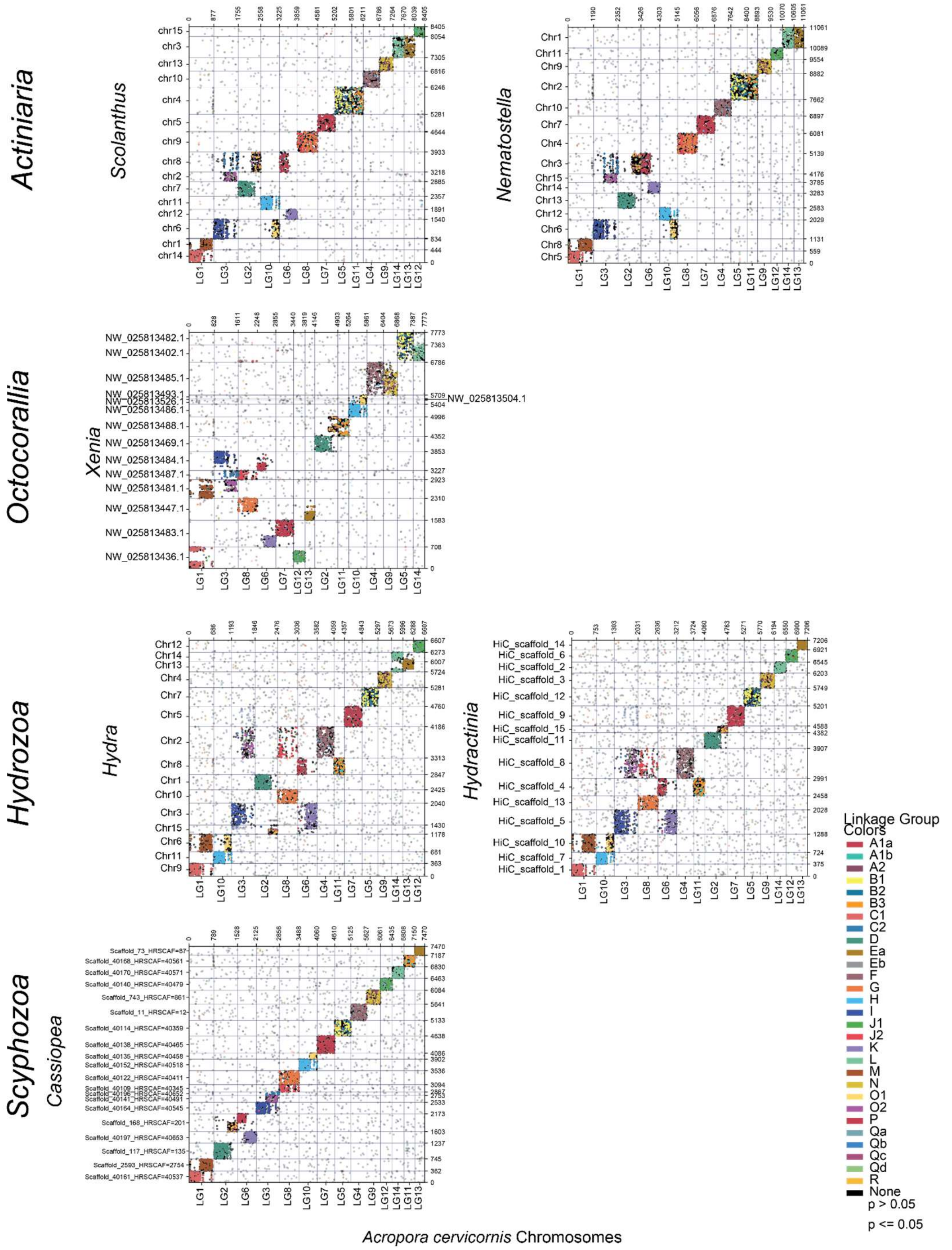


**Figure S3. Oxford dot plot of cnidarian assemblies against *A. cervicornis* chromosomes, arranged by taxonomic groups.** Ancestral linkage groups proposed for the last common ancestor of Cnidaria, Bilateria and sponges (Simakov et al. 2022) are indicated by color.

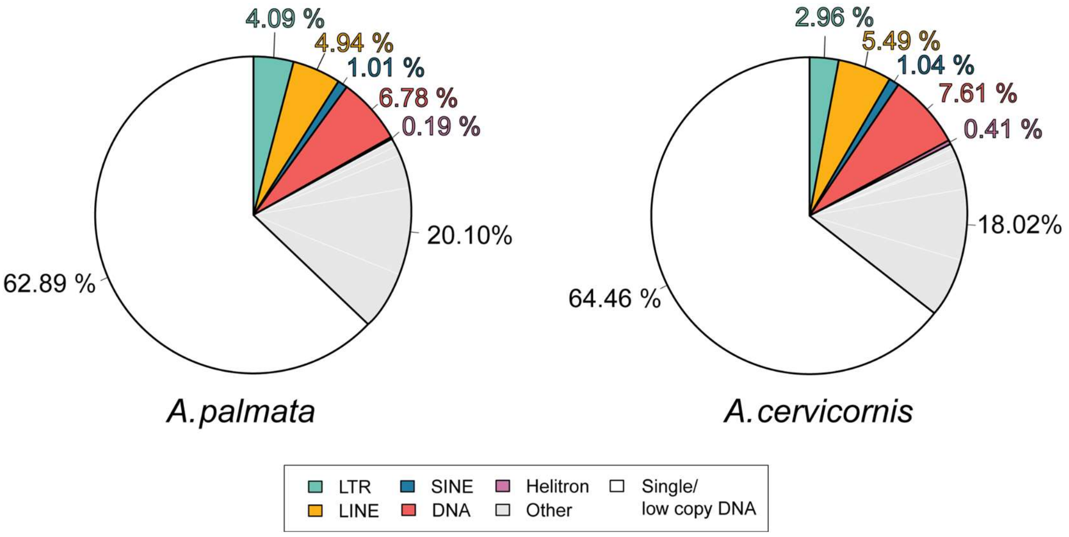


**Figure S4. Total repeat content estimates based on short-read sequence data of the Atlantic acroporids.** Repeat classes are colored as red = DNA, green = LTR, yellow = LINE, blue = SINE, purple = Helitron, and gray = Other (i.e., satellites, simple repeats, unannotated TEs).

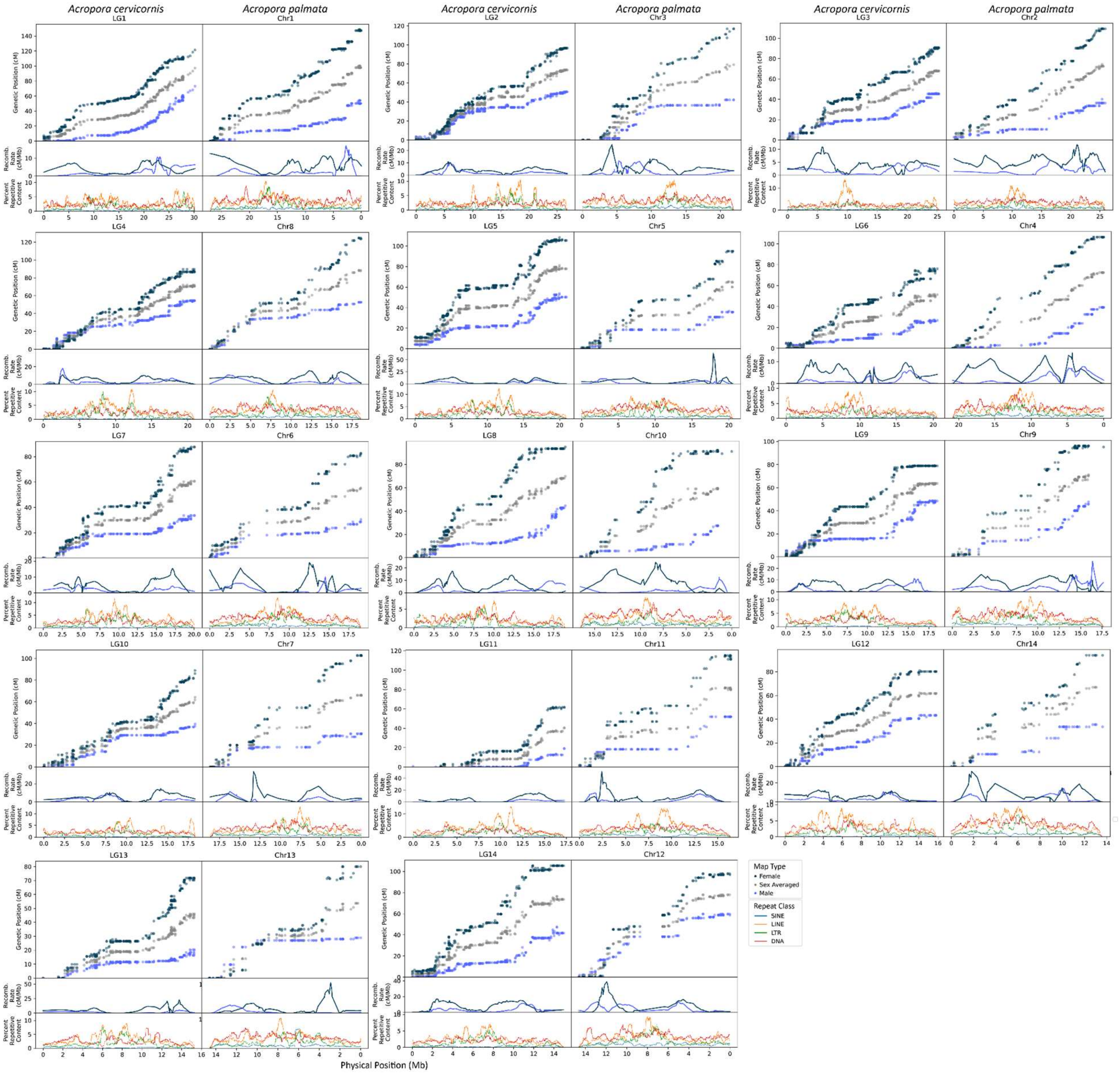


**Figure S5. Full linkage and recombination maps for all homologous chromosomes in the Atlantic acroporids.** LINE content appears to peak in regions with low recombination rates, highlighting centromere positions.

### Tables

**Table S1. Summary of sequencing libraries used for genome assembly of the Atlantic acroporids.**

| Species | Library Type | Tissue Type | Insert Size (bp) | Library Size (Gb) | Sequencer | Average Read Length (bp) | Read N50 (bp) | # of Reads (x $10^6$) | Genome Coverage | SRA Accession ID |
|---|---|---|---|---|---|---|---|---|---|---|
| *A. cervicornis* | Paired-end Illumina | Adult | 550 | 143.6 | HiSeq 2500 | 250 | | 313 | 489x | SRR7235995 |
| | Pacific Biosciences | Sperm | No Size Selection | 10 | Sequel | 3,238 | 4,394 | 3.1 | 25x* | |
| | Nanopore | Adult | No Size Selection | 94.4 | PromethION | 2,366 | 4,247 | 39 | 309x | |
| *A. palmata* | Paired-end Illumina | Sperm | 500 | 22 | HiSeq 2500 | 150 | | 74 | 69x | |
| | Paired-end Illumina | Adult | 550 | 114.6 | HiSeq 2500 | 250 | | 248 | 388x | SRR7235983 |
| | Pacific Biosciences | Sperm | 10,000 | 5.1 | Sequel | 7,126 | 10,110 | 0.71 | 13x* | |
| | Dovetail Chicago | Adult | NA | | Dovetail | 150 | | 200 | 165x | |
| | Dovetail Hi-C | Adult | N/A | | Dovetail | 150 | | 254 | 316x | |

* Estimated from Canu

**Table S2. Assembly statistics for the genome assemblies of the acroporids compared in this study.**

| | *A. palmata*[a] | *A. cervicornis*[a] | *A. digitifera*[b] | *A. hyacinthus*[c] | *A. millepora*[d] | *A. tenuis*[e] |
|---|---|---|---|---|---|---|
| # scaffolds | 406 | 3,308 | 955 | 907 | 854 | 614 |
| Assembly length (Mb) | 287.6 | 305.4 | 415.8 | 446.4 | 475.4 | 486.8 |
| Longest Scaffold (Mb) | 27.05 | 30.19 | 7.63 | 34.3 | 39.36 | 13.56 |
| Scaffold N50 (Mb) | 18.66 | 20.05 | 1.85 | 26.53 | 19.84 | 2.83 |
| Scaffold L50 | 7 | 7 | 69 | 8 | 9 | 46 |
| # gene models | 31,827 | 34,013 | 32,106 | 27,110 | 42,775 | 30,327 |
| BUSCO Completeness (Metazoa odb10) | C:87.8% [S:86.6%,D:1.2%],F:4.2%,M:8.0% | C:92.9% [S:92.6%,D:0.3%],F:3.6%,M:3.5% | C:92.2% [S:91.6%,D:0.6%],F:2.4%,M:5.4% | C:89.8% [S:89.4%,D:0.4%],F:2.5%,M:7.7% | C:93.4% [S:91.9%,D:1.5%],F:1.0%,M:5.6% | C:94.3% [S:93.1%,D:1.2%],F:1.3%,M:4.4% |

[a]Described in this study.
[b]NCBI GCA_014634065.1 genome assembly v2.0 (Shinzato et al. 2020)
[c]NCBI GCA_020536085.1 genome assembly (López-Nandam et al. 2023)
[d]NCBI GCF_013753865.1 genome assembly (Fuller et al. 2020)
[e]Reef Future Genomics 2020 Consortium (ReFuGe 2020 Consortium; Cooke et al. 2020 )

**Table S3 Assembly statistics showing completeness and contiguity of the *Acropora cervicornis* HG0582 (Selwyn and Vollmer 2023) and the HG0005 assembly (this study).** Mb = megabases. BUSCO = Benchmarking Universal Single-Copy Orthologs.

| | This Study | Selwyn and Vollmer 2023 |
|---|---|---|
| BUSCO Metazoa Complete Single Copy (%) | 92.80% | 92.03% |
| Complete Duplicated (%) | 0.30% | 0.42% |
| Fragmented (%) | 3.40% | 3.56% |
| Missing (%) | 3.50% | 3.98% |
| Assembly Size | 305.41Mb | 307.4Mb |
| Scaffold N50 | 20.051Mb | 2.8Mb |

**Table S4. K-mer genome size estimates.** Three different k-mer based genome size prediction tools were used: GenomeScope2, findGSE and kmergenie. The first two tools used the k-mer distribution histogram generated by Jellyfish with a k-mer size of 119. bp = base pairs.

| **Species** | **Jellyfish k119 + GenomeScope2** | **Jellyfish k119 + *find*GSE** | **kmergenie k119** |
|---|---:|---:|---:|
| *Acropora palmata* | 333,451,197 | 354,250,099 | 290,613,719 |
| *Acropora cervicornis* | 331,323,583 | 341,887,514 | 290,599,160 |

**Table S5. Comparison of *Acropora palmata* Hi-C (pseudo) chromosomes to *A. cervicornis* linkage groups. Mb = megabases.**

| *A. palmata* Chromosome IDs | *A. palmata* scaffold IDs | *A. palmata* Size (Mb) | *A. cervicornis* LG IDs | *A. cervicornis* Size (Mb) |
|---|---|---|---|---|
| Chr1 | hic_scaffold_4 | 27.05 | LG1 | 30.19 |
| Chr2 | hic_scaffold_10 | 25.92 | LG3 | 25.26 |
| Chr3 | hic_scaffold_2 | 21.90 | LG2 | 26.77 |
| Chr4 | hic_scaffold_20 | 20.87 | LG6 | 20.56 |
| Chr5 | hic_scaffold_17 | 20.54 | LG5 | 20.93 |
| Chr6 | hic_scaffold_31 | 19.01 | LG7 | 20.05 |
| Chr7 | hic_scaffold_35 | 18..66 | LG10 | 18.31 |
| Chr8 | hic_scaffold_15 | 18.58 | LG4 | 20.96 |
| Chr9 | hic_scaffold_6 | 17.67 | LG9 | 18.53 |
| Chr10 | hic_scaffold_30 | 16.55 | LG8 | 18.95 |
| Chr11 | hic_scaffold_5 | 16.42 | LG11 | 17.26 |
| Chr12 | hic_scaffold_1 | 14.66 | LG14 | 15.82 |
| Chr13 | hic_scaffold_21 | 14.61 | LG13 | 15.28 |
| Chr14 | hic_scaffold_11 | 13.63 | LG12 | 14.95 |

**Table S6. Number of structural variants identified between *Acropora palmata* and *A. cervicornis* genome assemblies. N/A = not applicable.**

| **Structural Variant Type** | **MUM&Co[1]** (O'Donnell and Fischer 2020) | **assemblytics** (Nattestad and Schatz 2016) | **SVIM-asm** (Heller and Vingron 2021) |
|---|---|---|---|
| Insertion* | 1004 | 3137 | 8448 |
| Deletion* | 610 | 2128 | 8014 |
| Tandem Duplication | 517 | 576 | 13 |
| Tandem Contractions | 209 | 199 | N/A |
| Interspersed Duplication | N/A | N/A | 3 |
| Repeat Expansions | 3843 | 1752 | N/A |
| Repeat Contractions | 3847 | 2908 | N/A |
| Inversions (<= 50kb and/or <2 marker support) | 155 | N/A | 27 |
| Translocations (<= 50kb and/or <2 marker support) | 165 | N/A | N/A |
| Inversions (=> 50kb, supported by at least 2 markers) | 5 | N/A | N/A |
| Translocations (=> 50kb, supported by at least 2 markers) | 177 | N/A | N/A |
| **TOTAL** | **10,532** | **10,700** | **17167** |

* Defined as "novel" insertion or deletion in MUMandCo.

**Table S7. Tracing the 21 cnidarian ancestral linkage groups as defined by Simakov et al. 2022 in acroporid corals. x = chromosomal fusion.** Acroporids share the same ALG architecture except for ALG L which is fused to G in *A. millepora* only.

| **Cnidarian ancestor** | ***Acroporids*** | | | | | | | | | | | |
|---|---|---|---|---|---|---|---|---|---|---|---|---|
| **21 ALGs** | ***A. hyacinthus*** | ***A. palmata*** | ***A. millepora*** | ***A. cervicornis* (LG #)** | **Same as Cnidarian Ancestor** | ***Catalaphyllia jardinei*** | ***Nematostella vectensis*** | ***Scolanthus sp.*** | ***Xenia sp.*** | ***Hydra vulgaris*** | ***Hydractinia symbiolongicarpus*** | ***Cassiopea xamachana*** |
| A1a | A1a | A1a | A1a | A1a (7) | Yes | A1a | A1a | A1a | A1axK | A1axC2 | A1aXC2 | A1a |
| RxQc | RxQcxD | RxQcxD | RxQcxD | RxQcxD (2) | No[1] | RxQcxD | | | | RxQcxKxI | RxQc | RxQcXP |
| D | | | | | | | D | D | D | | D | D |
| G | GxJ2 | GxJ2 | GxJ2 | GxJ2 (8) | No[2] | GxJ2 | GxJ2 | GxJ2 | GxEa | G | G | G |
| J2 | | | | | | | | | J2xC2 | | | J2 |
| H | HxO1 | HxO1 | HxO1 | HxO1 (10) | No[3] | HxO1 | H | H | H | H | H | H |
| O1 | | | | | | | O1xI | O1xI | | O1xM | O1xM | O1 |
| EbxFxQb | EbxF | EbxF | EbxF | EbxF (4) | No[4] | EbxF | EbxF | EbxF | EbxFxNxA2 | EbxFxJ2xO2xQd | EbxFxJ2xO2xQd | EbxFxQb |
| C1 | C1xM | C1xM | C1xM | C1xM (1) | No[5] | C1xM | C1 | C1 | | C1 | C1 | C1 |
| I | IxQdxO2xC2 | IxQdxO2xC2 | IxQdxO2xC2 | IxQdxO2xC2 (3) | No[6] | IxQdxO2xC2 | | | IxP | IxK | IxK | I |
| QdxO2 | | | | | | | QdxO2 | QdxO2 | MxO2xQd | | | QdxO2 |
| C2 | | | | | | | | | | | | C2 |
| K | KxP | KxP | KxP | KxP (6) | No[7] | KxP | K | K | | | | K |
| P | | | | | | | PxC2xRxQc | PxC2xRxQc | | | | |
| L | L | L | LxG | L (9) | Most[8] | L | LxEa | LxEa | LxB1xJ2xB2 | LxEa | L | L |
| NxA2 | NxA2 | NxA2 | NxA2 | NxA2 (14) | Yes | NxA2 | NxA2 | NxA2 | | NxA2 | NxA2 | NxA2 |
| Ea | Ea | Ea | Ea | Ea (13) | Yes | Ea | | | | | Ea | Ea |
| J1xQa | J1xQa | J1xQa | J1xQa | J1xQa (12) | Yes | J1xQa | J1xQa | J1xQa | C1xJ1xQa | J1xQa | J1xQa | J1xQa |
| B1xB2 | B1xB2 | B1xB2 | B1xB2 | B1xB2 (5) | Yes | B1xB2 | | | B1xB2 | B1xB2 | B1xB2 | B1xB2 |

| **Cnidarian ancestor** | ***Acroporids*** | | | | | | | | | | | |
|---|---|---|---|---|---|---|---|---|---|---|---|---|
| **21 ALGs** | ***A. hyacinthus*** | ***A. palmata*** | ***A. millepora*** | ***A. cervicornis* (LG #)** | **Same as Cnidarian Ancestor** | ***Cataphyllia jardinei*** | ***Nematostella vectensis*** | ***Scolanthus sp.*** | ***Xenia sp.*** | ***Hydra vulgaris*** | ***Hydractinia symbiolongicarpus*** | ***Cassiopea xamachana*** |
| A1bxB3 | A1bxB3 | A1bxB3 | A1bxB3 | A1bxB3 (11) | Yes | A1bxB3 | B1xB2xA1bxB3 | B1xB2xA1bxB3 | RxB3xA1bxQc | PxB3xA1b | PxB3xA1b | A1bxB3 |

[1]D fused to RxQc

[2]G and J2 fused

[3]H and O1 fused

[4]lost Qb

[5]C1 and M fused

[6]I and QdxO2 and C2 fused

[7]K and P fused

[8]part of G and L fused only in *A. millepora*

**Table S8. Genomic resources used to compare gene content and syntenic arrangement.** Genome assemblies and their predicted proteins were downloaded from sources noted below. Taxonomy is ranked as Subphylum, Class and Order.

| Species | Taxonomy | Source Type | Accession | Reference |
|---|---|---|---|---|
| *Acropora cervicornis* (genet GKR, HG0005) | Anthozoa, Hexacorallia, Scleractinia | This Study | | This Study |
| *Acropora cervicornis* (genet K2, HG0582) | Anthozoa, Hexacorallia, Scleractinia | NCBI | GCA_032359415.1 | Selwyn and Vollmer 2023 |
| *Acropora palmata* (genet HS1, HG0004) | Anthozoa, Hexacorallia, Scleractinia | This Study | | This Study |
| *Acropora tenuis* | Anthozoa, Hexacorallia, Scleractinia | OIST | aten_0.11.maker_post_001.proteins.fasta | Shinzato et al. 2021 |
| *Acropora digitifera* | Anthozoa, Hexacorallia, Scleractinia | NCBI | GCA_014634065.1 | Shinzato et al. 2011 |
| *Acropora millepora* | Anthozoa, Hexacorallia, Scleractinia | NCBI | GCF_013753865.1 | Fuller et al. 2020 |
| *Acropora hyacinthus* | Anthozoa, Hexacorallia, Scleractinia | NCBI | GCA_020536085.1 | López-Nandam et al. 2023 |
| *Montipora capitata* | Anthozoa, Hexacorallia, Scleractinia | USDA | Mcap_genemodels_1.1_aa.fas | Helmkampf et al. 2019 |
| *Orbicella faveolata* | Anthozoa, Hexacorallia, Scleractinia | NCBI | GCF_002042975.1 | Prada et al. 2016 |
| *Stylophora pistillata* | Anthozoa, Hexacorallia, Scleractinia | NCBI | GCF_002571385.2 | Voolstra et al. 2017 |
| *Catalaphyllia jardinei* | Anthozoa, Hexacorallia, Scleractinia | FigShare | Catalaphyllia_genomic.fasta | Yu et al. 2022 |
| *Amplexidiscus fenestrafer* | Anthozoa, Hexacorallia, Corallimorpharia | Reef Genomics | afen.prot.fa | Wang et al. 2017 |
| *Discosoma sp.* | Anthozoa, Hexacorallia, Corallimorpharia | Reef Genomics | dspp.prot.fa | Wang et al. 2017 |
| *Renilla muelleri* | Anthozoa, Octocorallia,Pennatulacea | Reef Genomics | renilla_augustus.aa | Jiang et al. 2019 |
| *Xenia sp.* | Anthozoa, Octocorallia,Alcyonacea | NCBI | GCF_021976095.1 | Hu et al. 2020 |
| *Exaiptasia diaphana* | Anthozoa, Hexacorallia, Actiniaria | NCBI | GCF_001417965.1 | Baumgarten et al. 2015 |
| *Nematostella vectensis* | Anthozoa, Hexacorallia, Actiniaria | NCBI | GCF_932526225.1 | Fletcher et al. 2023 |
| *Scolanthus callimorphus* | Anthozoa, Hexacorallia, Actiniaria | Simrbase | Scal100.fasta | Zimmermann et al. 2023 |
| *Cassiopea xamachana* | Medusozoa, Scyphozoa, Rhizostomeae | JGI | Casxa1_GeneCatalog_proteins_20200306.aa.fasta | Ohdera et al. 2022 |

| Species | Taxonomy | Source Type | Accession | Reference |
|---|---|---|---|---|
| *Hydra vulgaris* | Medusozoa, Hydrozoa, Anthoathecata | NCBI | GCF_022113875.1 | Simakov et al. 2022 |
| *Hydractinia symbiolongicarpus* | Medusozoa, Hydrozoa, Anthoathecata | NCBI | GCF_029227915.1 | Kon-Nanjo et al. 2023 |

**Table S9. Summary of repeat content in *Acropora palmata* and *A. cervicornis* genome assemblies.** Transposable elements highlighted in red are abundant among all acroporids.

| | | ***A. palmata*** | | | ***A. cervicornis*** | | |
|---|---|---|---|---|---|---|---|
| | | ***RepeatMasker*** | | ***dnaPipe TE*** | ***RepeatMasker*** | | ***dnaPipe TE*** |
| **Class** | | **Total bp Masked** | **% Masked** | **% Masked** | **Total bp Masked** | **% Masked** | **% Masked** |
| *Class I Retrotransposons* | | 21,084,824 | 6.93% | | 15,273,700 | 4.94% | |
| LTRs | TOTAL | 5,806,217 | 1.91% | 4.09% | 4,779,277 | 1.55% | 2.96% |
| | BEL | 47,737 | 0.02% | | | | |
| | Copia | 96,000 | 0.03% | | 53,260 | 0.02% | |
| | DIRS | 422,414 | 0.14% | | 548,673 | 0.18% | |
| | **Gypsy** | **3,622,045** | **1.19%** | | **27,631,128** | **0.89%** | |
| | Ngaro | 194,896 | 0.06% | | 110,327 | 0.04% | |
| | Pao | 1,423,125 | 0.47% | | 1,303,889 | 0.42% | |
| LINEs | TOTAL | 11,452,878 | 3.76% | 4.94% | 9,642,813 | 3.11% | 5.49% |
| | CR1 | 340,524 | 0.12% | | 429,999 | 0.14% | |
| | CRE | 338,153 | 0.11% | | 340,464 | 0.11% | |
| | Daphne | 384,411 | 0.13% | | | | |
| | Dong-R4 | 529,187 | 0.17% | | 333,832 | 0.11% | |
| | LINE1 | 1,117,505 | 0.37% | | 1,346,213 | 0.43% | |
| | LINE2 | 2,357,364 | 0.77% | | 2,305,268 | 0.74% | |
| | **Penelope** | **5,595,364** | **1.84%** | | **4,541,415** | **1.47%** | |
| | RTE | 324,903 | 0.11% | | 81,074 | 0.03% | |
| | Rex-Babar | 232,814 | 0.08% | | 231,723 | 0.07% | |
| | Others | 464,994 | 0.14% | | 32,825 | 0.01% | |

| Class | | *A. palmata* *RepeatMasker* Total bp Masked | % Masked | *dnaPipe TE* % Masked | *A. cervicornis* *RepeatMasker* Total bp Masked | % Masked | *dnaPipe TE* % Masked |
|---|---|---|---|---|---|---|---|
| *Class I Retrotransposons* | | 21,084,824 | 6.93% | | 15,273,700 | 4.94% | |
| LTRs | TOTAL | 5,806,217 | 1.91% | 4.09% | 4,779,277 | 1.55% | 2.96% |
| SINEs | TOTAL | 3,825,729 | 1.26% | 1.01% | 851,610 | 0.28% | 1.04% |
| | SINE1 | 145,314 | 0.05% | | | | |
| | SINE2 | 728,305 | 0.24% | | | | |
| | Others | 2,952,110 | 0.97% | | | | |
| *Class II DNA Transposons* | | 11,680,653 | 3.82% | 6.78% | 6,716,046 | 2.18% | 7.61% |
| | Academ | 803,610 | 0.26% | | 573,636 | 0.19% | |
| | CMC-EnSpm | 198,277 | 0.07% | | 217,306 | 0.07% | |
| | Crypton | 69,814 | 0.02% | | 28,423 | 0.01% | |
| | IS3EU | 31, 222 | 0.01% | | 38,852 | 0.01% | |
| | Kolobok | 117,329 | 0.04% | | 52,551 | 0.02% | |
| | MuDR | 229,628 | 0.07% | | | | |
| | Harbinger | 931,558 | 0.31% | | 525,479 | 0.17% | |
| | ISL2EU | 429,130 | 0.14% | | 434,799 | 0.14% | |
| | Polinton | 156,896 | 0.05% | | | | |
| | PiggyBac | 17,632 | 0.00% | | | | |
| | Sola | 784,761 | 0.26% | | 458,089 | 0.15% | |
| | **TcMariner** | **2,171,656** | **0.71%** | | **952,982** | **0.30%** | |

| Class | | *A. palmata* *RepeatMasker* Total bp Masked | *A. palmata* *RepeatMasker* % Masked | *A. palmata* *dnaPipe TE* % Masked | *A. cervicornis* *RepeatMasker* Total bp Masked | *A. cervicornis* *RepeatMasker* % Masked | *A. cervicornis* *dnaPipe TE* % Masked |
|---|---|---|---|---|---|---|---|
| *Class I Retrotransposons* | | 21,084,824 | 6.93% | | 15,273,700 | 4.94% | |
| LTRs | TOTAL | 5,806,217 | 1.91% | 4.09% | 4,779,277 | 1.55% | 2.96% |
| | **hAT** | **2,555,666** | **0.85%** | | **1,497,577** | **0.49%** | |
| | Others | 3,183,474 | 1.03% | | 1,988,903 | 0.65% | |
| RC- Helitron | | 358,513 | 0.12% | 0.19% | 79,119 | 0.03% | 0.41% |
| Unclassified TEs | | 17,561,493 | 5.77% | 14.87% | 32,033,663 | 10.35% | 13.26% |
| Low Complexity | | 10,424 | 0.00% | 0.63% | 488,815 | 0.16% | 1.00% |
| Simple Repeats | | 789,391 | 0.26% | 3.42% | 3,158,535 | 1.02% | 2.99% |
| Satellite | | 101,825 | 0.03% | 0.96% | 199,408 | 0.06% | 0.27% |
| rRNA | | 47,593 | 0.02% | 0.22% | 151,377 | 0.05% | 0.5% |
| snRNA | | 17,761 | 0.01% | | | | |
| **TOTAl** | | **50,778,858** | **16.69%** | **37.11%** | **58,510,908** | **18.91%** | **35.54%** |
